# Antisense therapy in a new rat model of Alexander disease reverses GFAP pathology, white matter deficits, and motor impairment

**DOI:** 10.1101/2021.01.29.428244

**Authors:** Tracy L. Hagemann, Berit Powers, Ni-Hsuan Lin, Ahmed F. Mohamed, Katerina L. Dague, Seth C. Hannah, Curt Mazur, Frank Rigo, Mel B. Feany, Ming-Der Perng, Robert F. Berman, Albee Messing

## Abstract

Alexander disease (AxD) is a devastating leukodystrophy caused by gain of function mutations in *GFAP*, and the only available treatments are supportive. Recent advances in antisense oligonucleotide (ASO) therapy have demonstrated that transcript targeting can be a successful strategy for human neurodegenerative diseases amenable to this approach. We have previously used mouse models of AxD to show that *Gfap*-targeted ASO suppresses protein accumulation and reverses pathology; however, the mice have a mild phenotype with no apparent leukodystrophy or overt clinical features and are therefore limited for assessing functional outcomes. In this report we introduce a new rat model of AxD that exhibits hallmark pathology with GFAP aggregation in the form of Rosenthal fibers, widespread astrogliosis, and white matter deficits. These animals develop normally during the first postnatal weeks but fail to thrive after weaning and develop severe motor deficits as they mature, with approximately 15 % dying of unknown cause between 6 to 12 weeks of age. In this model, a single treatment with *Gfap*-targeted ASO provides long lasting suppression, reverses GFAP pathology, and depending on age of treatment, prevents or mitigates white matter deficits and motor impairment. This is the first report of an animal model of AxD with myelin pathology and motor impairment, recapitulating prominent features of the human disease. We use this model to show that ASO therapy has the potential to not only prevent but also reverse many aspects of disease.

## INTRODUCTION

Alexander disease (AxD) is a rare neurogenetic disorder caused by dominant mutations in the gene encoding glial fibrillary acidic protein (*GFAP*), the major intermediate filament protein of astrocytes in the central nervous system. Age of onset ranges from infancy through adulthood, and symptoms include deficits in motor, cognitive, and brain stem functions, failure to thrive, and seizures. In most cases, the disease is progressive and fatal (*1*). The pathological hallmark of all forms of Alexander disease is the presence of astrocyte inclusion bodies known as Rosenthal fibers. Whether these protein aggregates are themselves toxic is not yet known. Nevertheless, what begins as primary dysfunction of astrocytes leads to a cascade of effects involving virtually every other cell type of the CNS (*2, 3*). Alexander disease has therefore become a unique model for primary astrocytopathy, with the potential to inform our understanding of astrocyte dysfunction in many neurological diseases.

As AxD is a monogenic gain-of-function disease, suppression of GFAP expression has recently emerged as the most promising therapeutic strategy (*4*). In mouse models engineered to carry *Gfap* point mutations mimicking disease-associated variants in humans, single intracerebroventricular (ICV) injections of *Gfap*-targeted antisense oligonucleotides (ASOs) reduced brain and spinal cord levels of GFAP transcript and protein to nearly undetectable levels and reversed key aspects of cellular and molecular pathology (*5*). However, the existing mouse models are not ideal. While they display several key features of the human disease, such as Rosenthal fibers, astrogliosis, and increased seizure susceptibility, they have no motor deficits, no evident leukodystrophy, and only subtle and strain-dependent deficits in cognition (*6, 7*). Hence the ability to fully evaluate the risks and benefits of GFAP suppression on clinically relevant phenotypes is limited.

Here we report a new rat model of AxD that displays more extensive pathology, including more severe behavioral symptoms and abnormalities of white matter. We find that administration of *Gfap*-targeted ASOs to these rats can prevent or reverse much of the pathology and symptoms, even when given at the peak expression of disease. These results offer new promise for the development of ASOs as an effective treatment strategy for an otherwise progressive and devastating disease.

## RESULTS

### Validation and phenotyping of GFAP mutant rat model

To generate a rat model of AxD, we used CRISPR/Cas9 mutagenesis to reproduce the severe R239H mutation observed solely in patients with early onset AxD at the orthologous GFAP-Arg237 position in the rat (Fig. 1A). Screening of founder animals identified two separate lines: one with the desired R237H mutation, and a second with a single nucleotide deletion upstream of the targeted sequence, generating a frameshift and premature stop codon (fig. S1). Sequence analysis of cloned cDNAs from rat brain in the R237H line demonstrates an intact full-length transcript from the targeted allele. Similar to patients with the disease and our mouse models (*7, 8*), expression of the R237H variant causes a spontaneous increase in *Gfap* transcript level (Fig 1B, left panel). In the second rat line, the deletion essentially creates a GFAP knockout rat with reduced transcript levels (Fig. 1B, right panel) and no detectable GFAP protein (Fig. 1C, right panel). We used this knockout line to cross with R237H rats to confirm protein expression from the targeted R237H allele (Fig. 1C, left panel). Immunoblotting with either a monoclonal antibody recognizing the GFAP tail domain or a polyclonal antibody demonstrates expression of the full length 50 kD protein. While the production of a GFAP null line provides a useful control, loss of GFAP function in the rat has no overt effects, similar to genetic knockout in the mouse (*9–12*). Since AxD associated point mutations lead to a toxic gain of function for the encoded protein (*13*), we focus here on the heterozygous point mutant R237H rat.

**Fig. 1.**
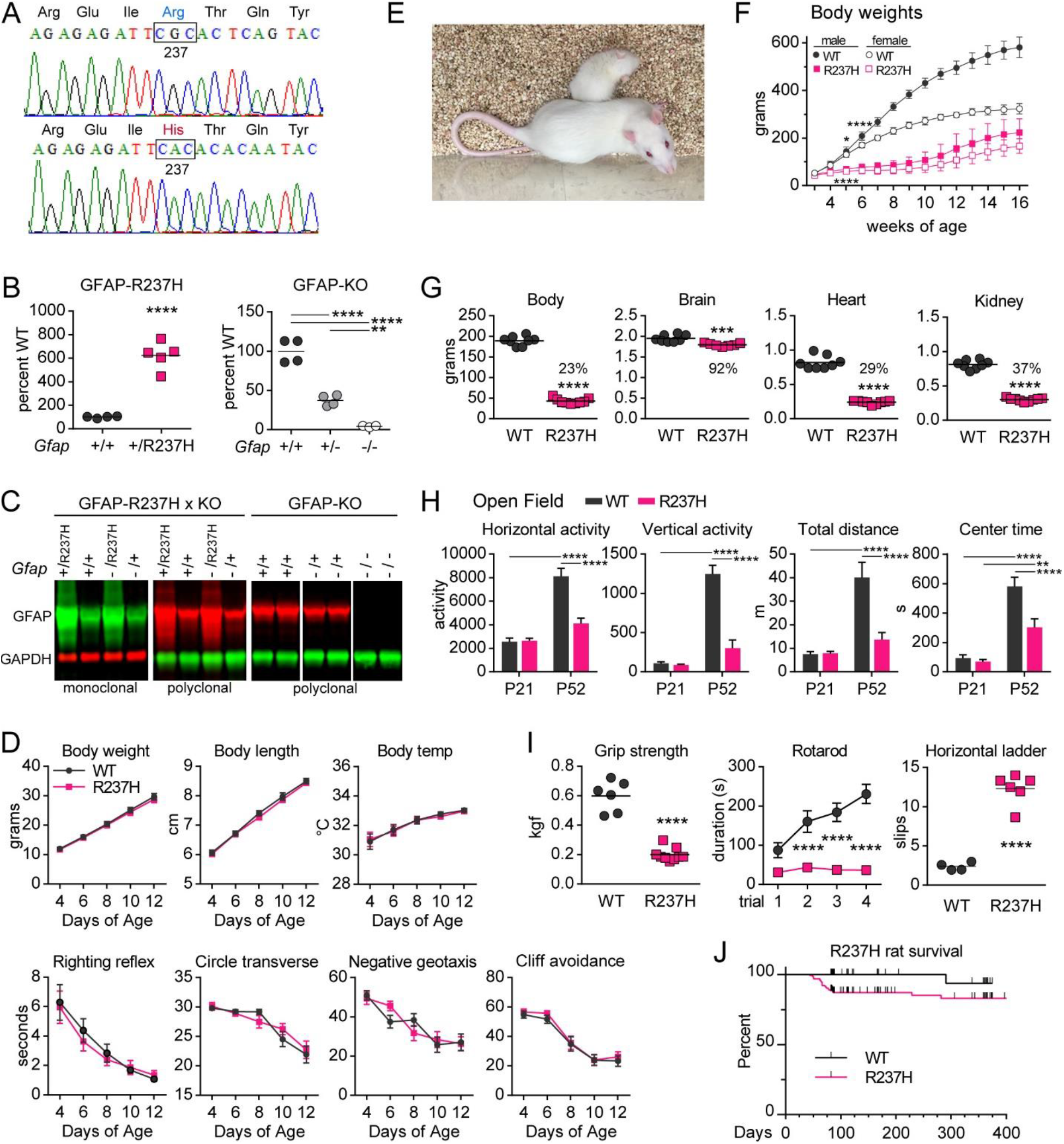
Validation and phenotyping of GFAP mutant rat model. (**A**) Sequence analysis of cDNA from R237H rat brain shows successful CRISPR/Cas9 targeting for AxD-associated missense mutation. (**B**) Quantitative PCR demonstrates increased *Gfap* transcript in R237H rat brain at postnatal day 21 (P21, left panel, ****p < 0.0001, 2-tailed t-test), while transcript in a second line harboring a *Gfap* frameshift mutation shows incremental decreases in *Gfap* in heterozygous and homozygous rat brain (P30, right panel, ****p<0.0001, **p=0.0069, 1-way ANOVA, Tukey’s multiple comparisons). (**C**) Western analysis of total protein from GFAP point mutant (R237H) and knockout (KO) rats demonstrates increased levels of full-length protein in the heterozygous mutants and loss of protein in the knockout with both monoclonal (GA5) and polyclonal (DAKO Z0334) antibodies. Crosses between the two lines to combine mutant and null alleles (*Gfap* −/R237H) confirms expression of the R237H targeted allele. (**D**) R237H rats meet normal developmental milestones during the second postnatal week (no effect of genotype, 2-way repeated measures ANOVA, N = 15 litters consisting of 39 WT (20 male, 19 female) and 39 R237H (19 male, 20 female), error bars = SEM). (**E** and **F**) By 5 weeks of age, R237H rats show significant differences in body weight compared to their wild-type littermates and by 8 weeks are less than one third the size (males shown in **E**, 2-way repeated measures ANOVA, Sidak’s multiple comparisons show significant differences by 5 weeks of age, *p = 0.0138, males, ****p < 0.0001, females, weeks 6-13, p < 0.0001 for males and females, N = 4-6, error = standard deviation). (**G**) Organ weights are also reduced, although brain is not as severely affected (percent of WT shown above R237H, ***p = 0.0006, ****p < 0.0001, 2-tailed t-test, females at 10 weeks). (**H**) Open field measures show normal activity of R237H rats at P21, but significant differences in horizontal and vertical movement, total distance traveled (m = meters) and time in the center of the arena (s = seconds) at P52 compared to WT (two-way ANOVA, Tukey’s multiple comparisons, N = 10 (5 males, 5 females) per genotype, error bars = SEM). (**I**) Measures of forelimb grip strength (****p < 0.0001, 2-tailed t-test, females at 8 weeks), rotarod performance (2-way repeated measures ANOVA, Sidak’s multiple comparisons show differences in trials 2-4, ****p < 0.0001, males and females at 8 weeks, N = 9-11, error bars = SEM) and average number of slips per run on a horizontal ladder (****p < 0.0001, 2-tailed t-test, females at 8 weeks), demonstrate motor deficits in R237H rats. (**J**) Kaplan-Meyer curve of survival rates of R237H rats aged to 12 weeks or more shows 14 % die or are found moribund and euthanized (p = 0.0008, Mantel-Cox test, N = 165 R237H, 98 WT male and female rats). Black tick marks indicate animals removed for experiments.

R237H rats meet normal developmental milestones (Fig. 1D) and are physically difficult to distinguish from their wild-type littermates during the first 3 postnatal weeks. However, after weaning both males and females fail to thrive and by 8 weeks are gaunt, with virtually no white fat (fig. S2) and reduced organ size (Fig. 1, E to G), although brain is relatively spared. These differences are further exemplified by open field activity where weanlings show no difference between genotypes, but the same animals at 8 weeks show marked reduction in activity (Fig 1H). Adult R237H rats have reduced strength, deficits in coordination, and gait abnormalities, as demonstrated by forelimb grip, rotarod performance, and paw placement on a horizontal ladder (Fig. 1I, Movies S1 and S2). While genotype and sex ratios are normal at weaning (R237H: 24.4% male, 24.8 % female; wildtype: 24.6 % male, 26.2 % female; N=1,428), approximately 14 % of AxD rats die between 6-12 weeks (including 5.5% found moribund or with severe hemi/paraparesis or paralysis and euthanized for humane reasons) (Fig. 1J). Occasional surviving rats develop forelimb monoplegia between 6 to 8 weeks of age.

### GFAP accumulation, Rosenthal fibers, and astrogliosis in R237H rats

*GFAP* mutations in AxD lead to GFAP accumulation, protein aggregation in the form of Rosenthal fibers, and astrogliosis. To determine whether R237H rats replicate key pathogenic features of human disease, we first assessed GFAP levels and protein aggregation as the animals mature. By three weeks of age, brain GFAP is significantly increased and by 8 weeks dramatically elevated (Fig. 2A). In mutant animals, lower molecular weight fragments are also apparent with immunoblotting and most likely reflect 24 and 26 kD caspase-6 cleavage products (*14, 15*). An accumulation of high molecular weight ubiquitinated proteins is also apparent (Fig. 2B), suggesting diminished proteasome capacity. Further analysis of Rosenthal fiber-associated proteins shows an increase in small heat shock proteins Hsp27 (HSPB1) and αB-crystallin (CRYAB), autophagy cargo protein sequestosome (p62, SQSTM1), intermediate filament vimentin (VIM), and a more recently identified component, cyclinD2 (CCND2, Fig. 2B) (*16–19*).

**Fig. 2.**
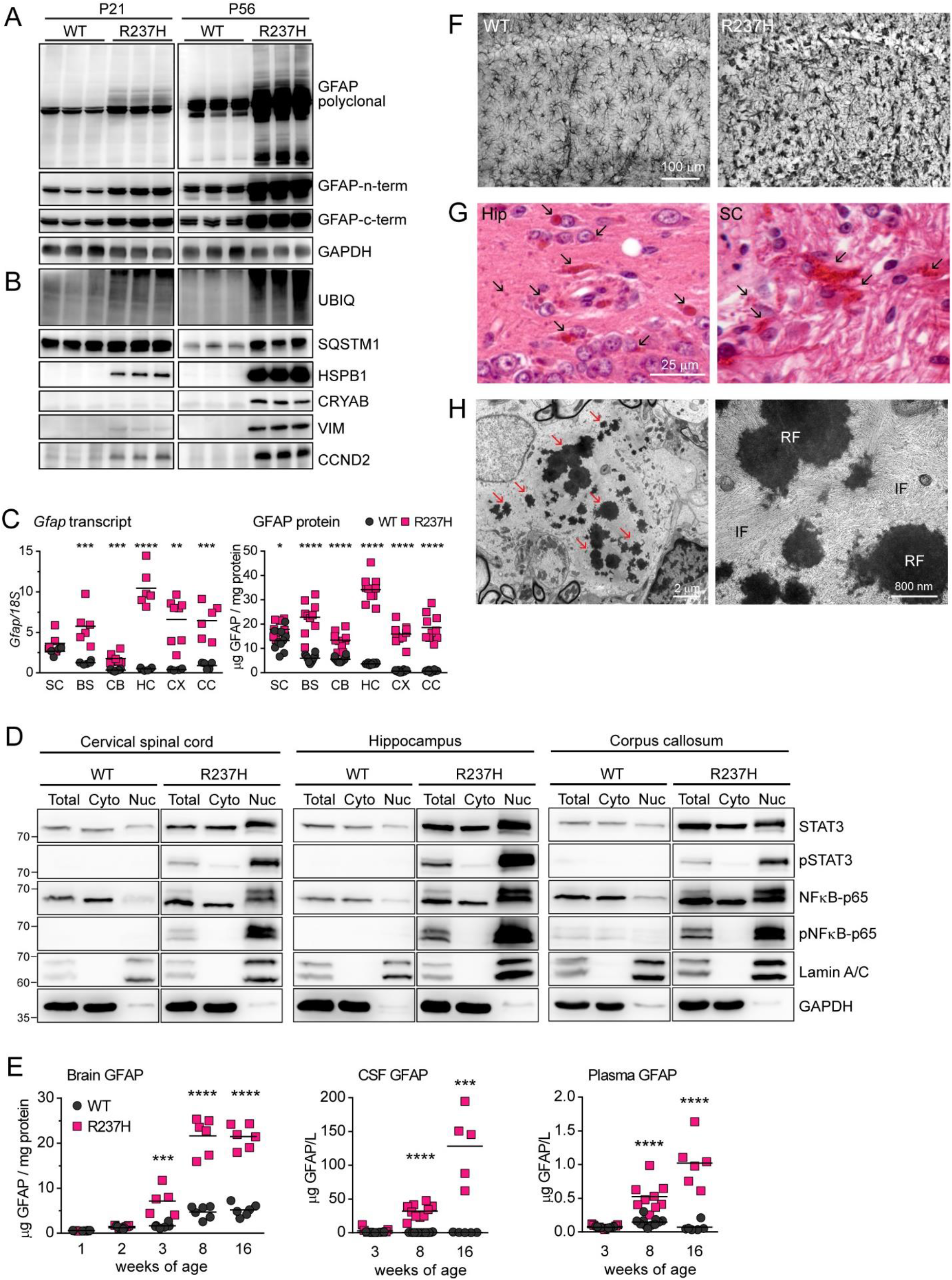
GFAP accumulation, Rosenthal fibers, and astrogliosis in R237H rats. (**A**) Western analysis of GFAP protein at P21 and P56 shows early GFAP accumulation in R237H rat brain. Three different antibodies targeting various epitopes are used to verify the presence and size of GFAP from mutant animals. (**B**) Western analysis of proteins associated with Rosenthal fibers and astrocyte pathology in AxD shows increases in stress proteins SQSTM1, HSPB1, CRYAB, and VIM as well as CCND2. Lanes represent individual animals. (**C**) Regional transcript (qPCR) and protein (ELISA) analysis at 8 weeks of age (both sexes) shows significant increases in GFAP over wild-type animals in all brain regions analyzed, with the exception of spinal cord transcript (*p < 0.05, **p < 0.01, ***p < 0.001, ****p < 0.0001, multiple t-tests, Holm-Sidak correction method, α = 5 %, N = 4-8 in C, N = 10 in D). (**D**) Western analysis of total protein and subcellular lysate fractions from cervical spinal cord, hippocampus and corpus callosum show increases of phosphorylated STAT3 and NFκB in the nuclear compartment (Nuc) of R237H rat CNS. Lamin A/C is used as a nuclear protein control, but also shows an increase in R237H rats. GAPDH predominantly localizes in the cytosolic fraction (Cyto), N = 3, representative blots shown. (**E**) GFAP-ELISA of brain lysates, CSF, and plasma show significant increases in GFAP levels as R237H rats age (***p < 0.001, ****p < 0.0001, multiple t-tests, Holm-Sidak correction method, α = 5 %, N = 5-6 for brain, N = 6-11 for plasma, N = 5-14 for CSF). (**F**) Immunohistochemistry demonstrates astrogliosis with GFAP accumulation (hippocampus, CA1 shown at 8 weeks of age). (**G**) H&E stain shows Rosenthal fibers as eosinophilic inclusions and puncta (arrows, not all indicated) in hippocampus and spinal cord at 8 weeks of age. (**H**) Electron microscopy shows hallmark Rosenthal fibers as dense osmiophilic aggregates ensnared in a bed of intermediate filaments (arrows on left panel indicate representative fibers; enlarged in right panel, RF = Rosenthal fiber, IF = intermediate filaments).

In mouse models of AxD, astrocyte pathology is localized and more prominent in forebrain structures, with limited pathology in hindbrain (*7*). GFAP expression in the adult R237H rat is markedly elevated, at both the transcript and protein level, in all regions analyzed including brainstem, cerebellum, hippocampus, cortex and corpus callosum, with the exception of spinal cord (which begins with the highest basal expression) (Fig. 2C, 8 weeks of age). While an inadequate response by the ubiquitin-proteasome system likely exacerbates protein accumulation, increased levels of GFAP in AxD are also driven at the transcriptional level (*5, 20, 21*). Western analysis of subcellular protein fractions indicates activation of both STAT3 and NFκB, transcription factors known to regulate *Gfap*, as evidenced by elevation of both native and phosphorylated forms of the two, and localization of phosphorylated proteins in the nuclear compartment (Fig. 2D). Immunofluorescent labeling confirms nuclear localization of phosphorylated STAT3 in GFAP positive astrocytes and potentially other cell types (fig. S3). Note that the nuclear membrane intermediate filament lamin A/C, used in these experiments to confirm enrichment of nuclear proteins (Fig. 2D), is also elevated in the R237H rat. These results agree with our previous report showing elevation of A-type lamin in both animal models and patients with AxD, implicating mechanotransduction signaling in AxD pathogenesis (*22*). GFAP levels in CSF and plasma also increase as AxD rats mature, with significant elevation by 8 weeks of age, when brain levels are maximal and animals are severely affected (Fig. 2E). Elevation of GFAP in these two body fluid compartments may provide useful biomarkers for therapeutic testing in the model.

Hypertrophic reactive astrocytes (*23*) are apparent throughout the R237H rat CNS and have abnormal morphology with protein aggregates and thick blunted processes (Fig. 2F, fig. S4). Rosenthal fibers are evident by light microscopy as early as P14 and are prominent as both large aggregates and smaller puncta in brain and spinal cord of young adult animals (Fig. 2G, fig. S5). Importantly, electron microscopy demonstrates classic Rosenthal fibers as observed in astrocytes of patients with AxD: dense osmiophilic aggregates enmeshed in swirls of intermediate filaments (Fig. 2H).

### Astrocyte stress response in R237H rats: pathology and function

Although cellular GFAP is predominantly filamentous, the protein normally exists in equilibrium between a detergent-soluble pool of monomers, dimers, and tetramers and an insoluble pool of unit length and full filaments. Biochemically, Rosenthal fibers in AxD are even less soluble and can be partitioned further from the filaments (*19*). In R237H rats, total GFAP protein is elevated throughout the brain, but in spinal cord, where GFAP levels are normally high, no further elevation is observed (Fig. 2C). To determine whether GFAP aggregates have similar properties in the R237H rat and if brain and spinal cord show differences in these pools, we separated protein lysates from hippocampus and cervical cord into soluble, cytoskeletal, and Rosenthal fiber (RF) enriched fractions (*19*). In hippocampus, GFAP was elevated in all three pools, while in spinal cord GFAP showed little change in the cytoskeletal fraction, but a marked increase in the RF fraction (Fig. 3, A and B). These data show that the presence of mutant GFAP by itself, independent of GFAP elevation, in the spinal cord is sufficient to produce AxD-associated pathology.

**Fig. 3.**
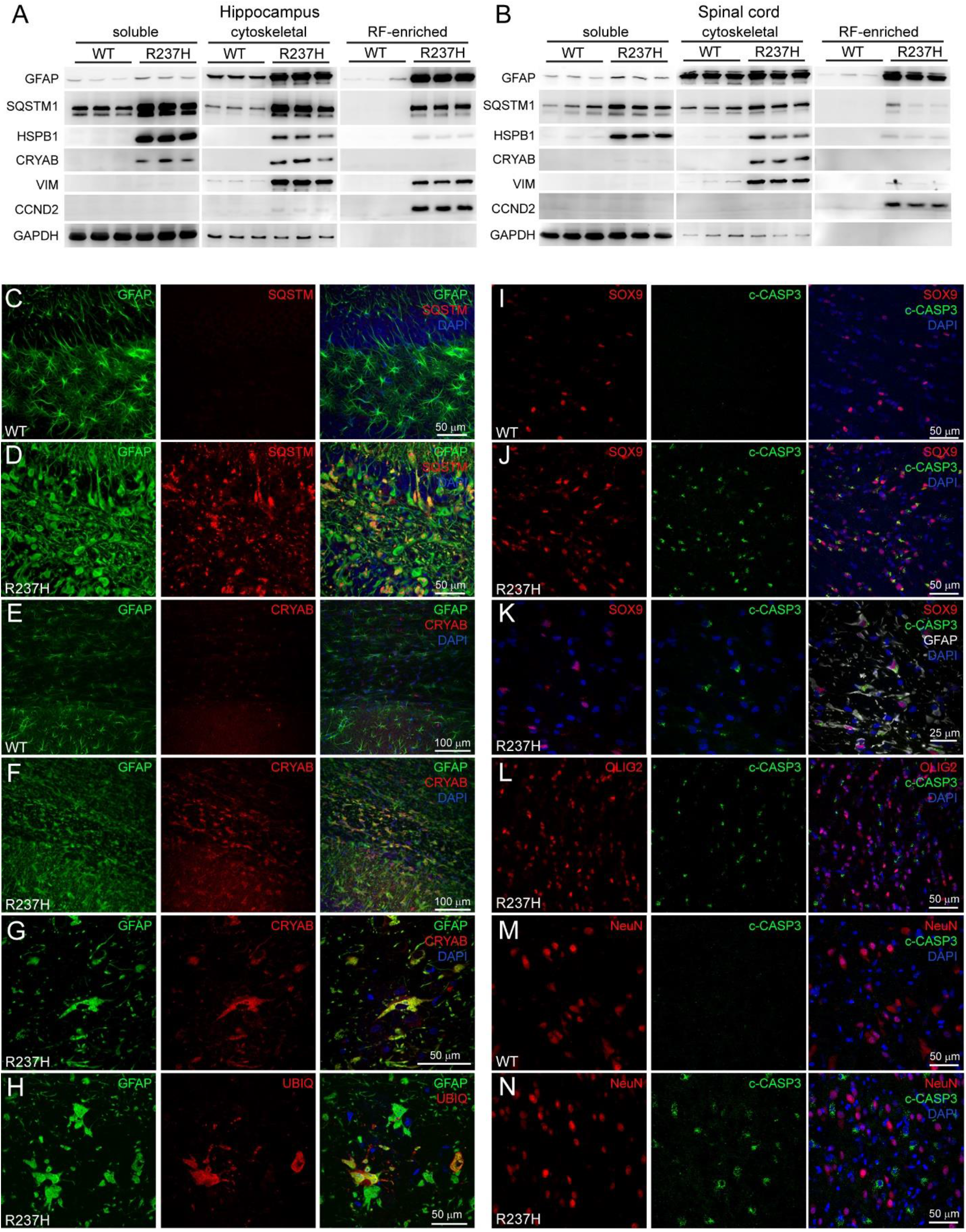
Astrocyte stress response and pathology in R237H rats. (**A** and **B**) Western analysis of soluble, cytoskeletal, and Rosenthal fiber (RF) enriched fractions of GFAP in hippocampus (**A**) and cervical spinal cord (**B**) shows marked accumulation of GFAP in the RF-enriched fraction in R237H rats. Other RF associated proteins show different patterns of accumulation among the different fractions (N=3, with each lane representing individual animals). (**C** and **D**) Immunofluorescent labeling (IF) shows SQSTM1 is elevated and colocalizes with GFAP aggregates in R237H rats (dentate gyrus shown). (**E** to **G**) CRYAB-IF shows a similar pattern in corpus callosum (**F** and **G**), in contrast to wild-type animals (**E**) where CRYAB is typically expressed by oligodendrocytes. (**H**) UBIQ-IF can be found in astrocytes, but does not completely colocalize with GFAP (hippocampal fissure). (**I**to N) Cleaved caspase-3 positive astrocytes are evident in corpus callosum (shown, **I** to **K**) and other brain regions as shown by colocalization with Sox9 and GFAP, but other cell types including those of the oligodendrocyte lineage (**L**, Olig2, corpus callosum) and neurons (**M** and **N**, NeuN, brainstem) do not show evidence of caspase-3 activation in R237H rats at 8 weeks of age. Confocal images are single optical slices, N = 5 animals per genotype (males and females). Camera and microscope settings were equivalent for comparisons between genotypes.

We also analyzed AxD-associated stress response proteins. SQSTM1 displayed a pattern similar to GFAP in both CNS regions, but other stress proteins varied in elevation and cellular compartment. HSPB1 was elevated in all fractions in both regions, while CRYAB was surprisingly not detectable in the RF fraction. CCND2 was predominantly detected in the RF fraction. Note that most cytoplasmic proteins are extracted in the soluble fraction, including GAPDH. Histologically, immunofluorescent labeling of SQSTM1 showed colocalization with GFAP and provides a robust marker for GFAP protein aggregation (Fig. 3, C and D). CRYAB, which is normally primarily expressed in oligodendrocytes, is elevated in R237H rat astrocytes and appears to colocalize with GFAP (Fig. 3, E to G). Ubiquitin labeling was less prevalent and did not always colocalize with GFAP, which may reflect accumulation of other ubiquitinated proteins (Fig. 3H).

Prior studies of astrocyte response to injury and disease, including AxD, have implicated caspase-3 activation (*24, 25*). To evaluate astrocyte injury, we examined whether cleaved caspase-3 was present in the R237H rat. We found prominent caspase-3 activation apparent in hippocampus, corpus callosum and other regions of the CNS as early as 3 weeks of age when the overall clinical presentation is still relatively mild (fig. S6). Co-labeling with markers for astrocytes (Sox9, GFAP), oligodendrocytes (Olig2), and neurons (NeuN) in adult animals showed that cleaved caspase-3 was specifically localized to astrocytes (Fig. 3, I to N). Further analysis with TUNEL identified a small population of apoptotic cells, and while there were more TUNEL positive cells in R237H rats compared to wild-type, not all positive cells could be identified as astrocytes (fig. S6D). Caspase-3 activation has been associated with reactive astrocytes in other injury models, particularly in rats, and may have non-apoptotic functions such as cytoskeletal rearrangement (*26–30*).

To evaluate potential effects of GFAP induced pathology on astrocyte function, we analyzed the glutamate transporter GLT-1 (SLC1A2), the major gap junction protein in astrocytes, connexin 43 (GJA1), water channel aquaporin 4 (AQP4), and potassium channel Kir4.1 (KCNJ10). R237H rats show a dramatic loss of SLC1A2 by 8 weeks of age as shown by immunoblotting of membrane fractions from different brain regions and spinal cord (Fig. 4A, total lysates in fig. S7), consistent with reductions observed in mouse models and patients with AxD (*31*). Previous studies with in vitro models have also suggested a lack of gap junction connectivity in astrocytes expressing mutant GFAP (*31*). Immunofluorescent labeling of GJA1 in hippocampus shows prominent expression in the hilus and near the fissure vasculature in wild-type animals, but patchy labeling with more puncta located away from vessels in R237H rats (Fig. 4, B and C). In spinal cord GJA1 expression is generally reduced (Fig. 4, D and E), and membrane tissue fractions show a decrease in GJA1 levels in all regions analyzed (Fig. 4A). AQP4 distribution also appears to be altered and depolarized away from astrocyte endfeet in hippocampus, particularly in the hilus, and is reduced in spinal cord (Fig. 4, A, F to I). KCNJ10 which tends to distribute in the same subcellular compartments as AQP4 (*32*), is localized to astrocyte somata in hippocampus and virtually absent in spinal cord (Fig. 4, A, J to M). AQP4, KCNJ10 and GJA1 normally localize to astrocyte endfeet and regulate ion and water transport. To determine whether edema may contribute to CNS pathology in the R237H rat, we measured water content in different brain regions and spinal cord. With the exception of cerebellum, we found significant increases in all regions tested (Fig. 4N), with the most dramatic changes occurring in areas predominantly composed of white matter including spinal cord, brainstem and corpus callosum. A small decrease was observed in cerebellum. To assess blood brain barrier integrity, we measured IgG content in hippocampus and spinal cord and found modest elevation in hippocampus (Fig. 4O).

**Fig. 4.**
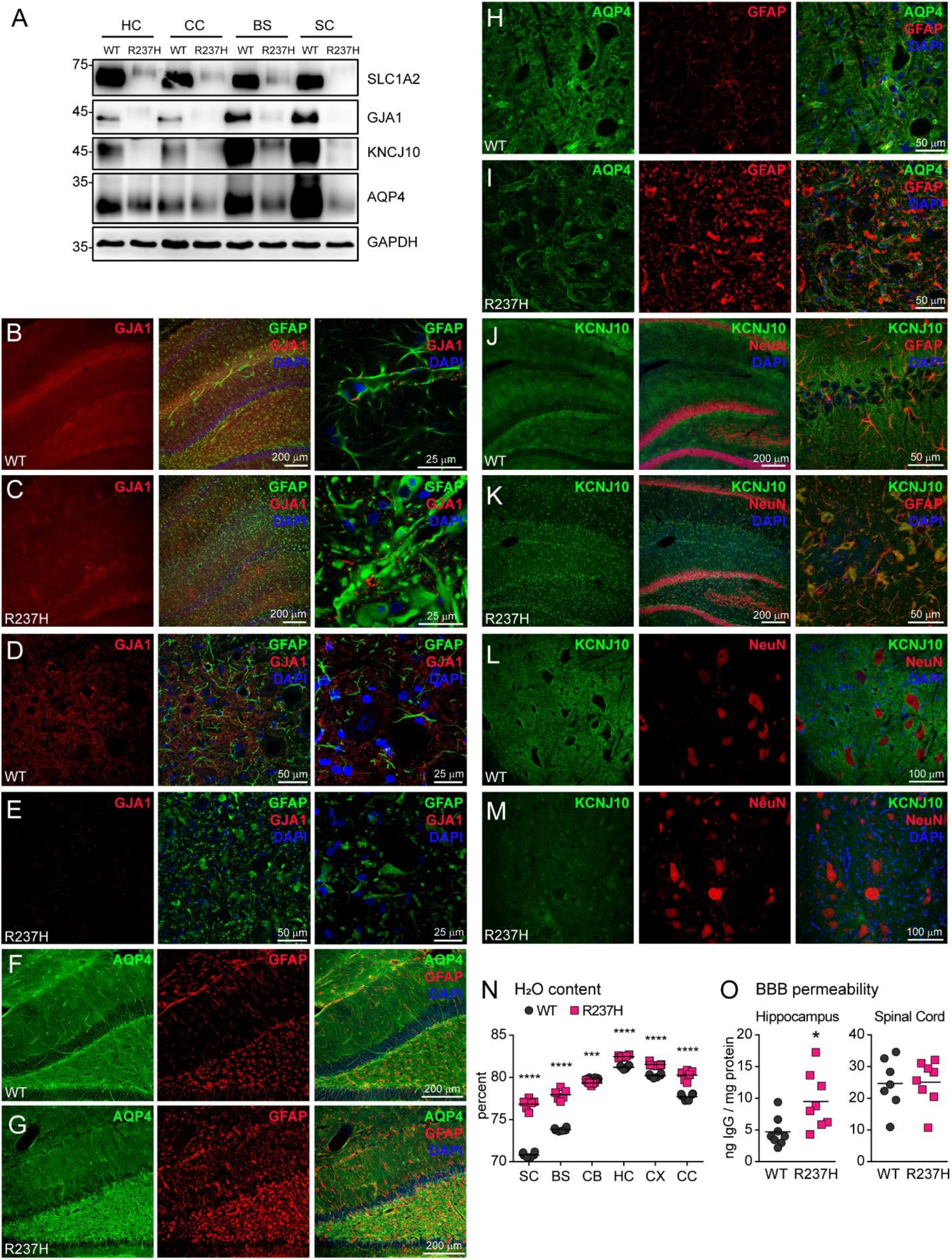
Astrocyte function in R237H rats. (**A**) Immunoblotting of membrane fractions from hippocampus (HC) corpus callosum (CC), brainstem (BS) and cervical spinal cord (SC) shows changes in proteins related to astrocyte function including glutamate transporter GLT1 (SLC1A2), connexin 43 (GJA1), aquaporin 4 (AQP4) and Kir4.1 (KCNJ10) in R237H rats compared to WT at 8 weeks of age (GAPDH loading control, N = 4 males, representative lysates shown). (**B** to **E**) Immunofluorescence (IF) shows altered distribution of GJA1 in hippocampus (**B** and **C**), and in spinal cord (cervical ventral horn), labeling is generally less prominent (**D** and **E**). (**F** to **I**) AQP4-IF shows depolarization in hippocampus (**F** and **G**) and reduction in spinal cord (**H** and **I,** cervical ventral horn). (**J** to **M**) KCNJ10 is mislocalized to astrocyte cell bodies in hippocampus (**J** and **K,** left panels show low magnification with NeuN labeling for orientation and right panel shows higher magnification with GFAP co-label) and IF shows a marked decrease in spinal cord gray matter (**L** and **M**). Confocal images are single optical slices, with the exception of higher magnification images in D, E, J, K (right panels), which are maximum projections. Camera and microscope settings were equivalent for comparisons between genotypes. N = 5 animals per genotype (males and females), and images are representative. (**N**) Water content in different CNS regions shows significant increases in all but cerebellum in R237H rats. Water content in cerebellum is decreased. (***p < 0.001, **** p <0.0001, multiple t-tests, Holm-Sidak correction method, α = 5 %, N = 6 males). (**O**) Quantification of IgG content by ELISA to assess blood brain barrier integrity in saline perfused female rats at 10 weeks of age shows no change in spinal cord but a modest increase in hippocampus (*p < 0.05, 2-tailed t-test, N = 8). All results from rats at 8 weeks except for **O**. SC = cervical spinal cord, BS = brainstem, CB = cerebellum, HC = hippocampus, CX = cortex, CC = corpus callosum.

### Non-cell-autonomous effects of mutant GFAP and astrocyte dysfunction

Given the molecular and morphological changes in AxD astrocytes and the activation of neuroinflammatory pathways, we wanted to survey how these changes affect other cell types in the R237H rat CNS. Not surprisingly, Iba1 immunolabeling shows an increase in hypertrophic microglia in most regions of the CNS. In areas such as cortex and spinal cord gray matter (Fig. 5, A to E), microglia appear to wrap around GFAP-laden astrocytes (Fig. 5, B and E), with some having a reactive ameboid morphology (Fig. 5D). Oligodendrocyte precursor cells show an increase in NG2 immunolabeling particularly in white matter (Fig. 5, F and G), suggesting this glial population is also responding to astrocyte pathology. Identification of mature oligodendrocytes with APC/CC1 or glutathione S-transferase (GST)-π was unfortunately hampered by cross reactivity with reactive astrocytes (*33–35*).

**Fig. 5.**
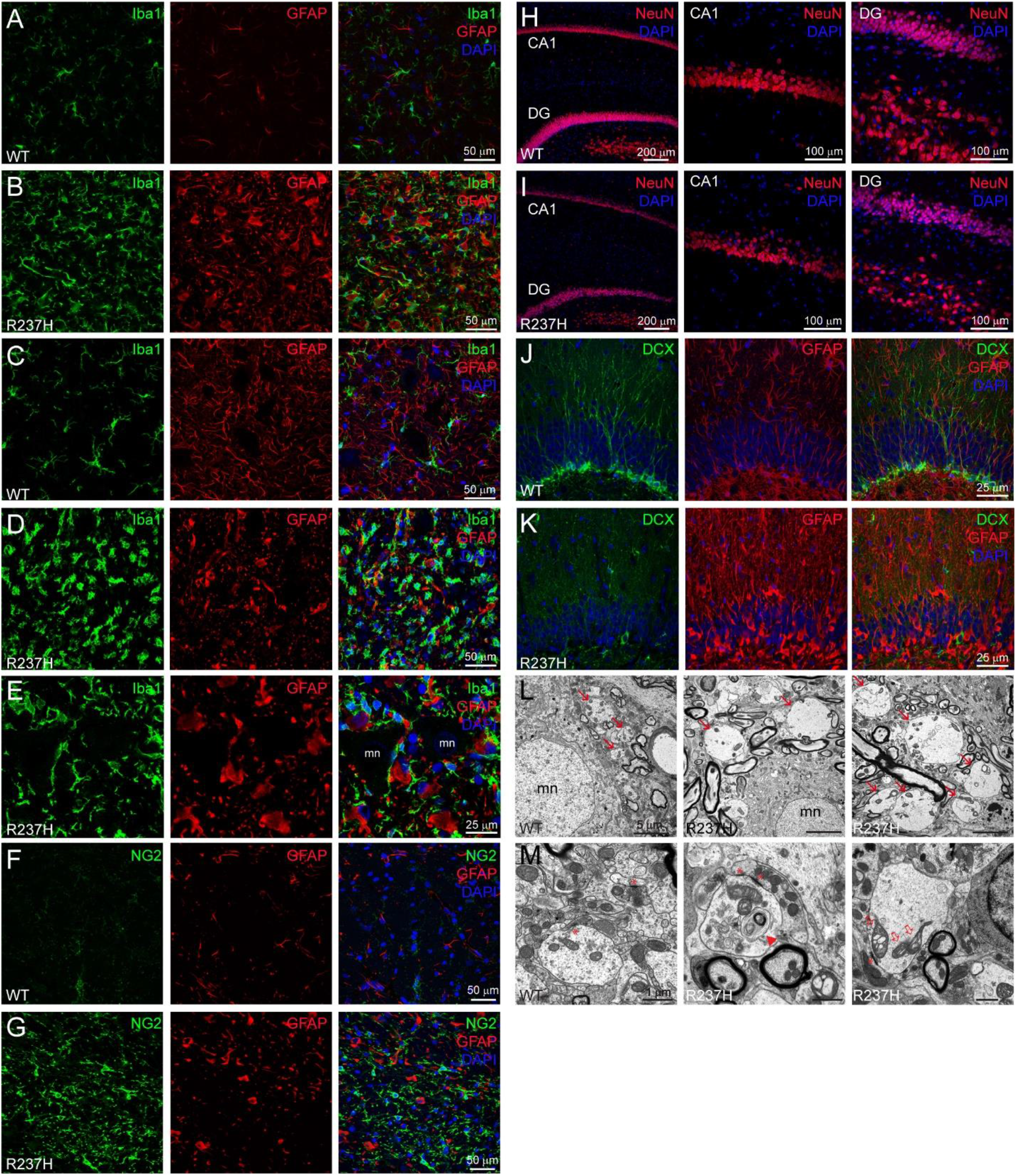
Non-cell-autonomous effects of mutant GFAP and astrocyte dysfunction. (**A** to **E**) Microglia identified by Iba1 immunofluorescence (IF) are hypertrophied in R237H rats (**B**, **D**, and **E**) and in some cases ameboid (particularly in **D**) in most regions of the CNS (cortex in **A** and **B,**cervical cord in **C** to **E**) and often found in intimate association with astrocytes in R237H rats at 8 weeks of age (mn = motoneuron in **E**). (**F** and **G**) NG2-IF shows reactive oligodendrocyte precursor cells particularly in white matter regions (corpus callosum shown). (**H** and **I**) NeuN-IF highlights thinning pyramidal cell (CA1) and dentate granule cell (DG) layers in R237H rats. (**J**and **K**) Doublecortin (DCX) IF shows a lack of new neurons in dentate gyrus reflecting deficits in adult neurogenesis is R237H rats. (**L** and **M**) Electron microscopy in cervical cord ventral horn from R237H rats shows enlarged neurites (**L**, arrows), some of which show degeneration and collections of membranous debris (**M**, arrow head) and vacuolized mitochondria (open arrows). Asterisks indicate synaptic densities.

Within the dense collection of reactive glia, neurons show signs of degeneration and loss. In hippocampus, both the pyramidal and the dentate granule cell layers appear mottled and thinner, suggesting a reduced neuronal population (Fig 5, H and I). We have previously reported that GFAP mutations lead to deficits in adult neurogenesis in mouse models of AxD (*6*), and R237H rats have virtually no doublecortin (DCX) positive neurons in the dentate gyrus at 8 weeks of age (Fig. 5, J and K). Whether the lack of new neurons is the result of deficits in the adult radial glia-like stem cells, which express GFAP, or a reactive subgranular zone niche remains to be determined. In spinal cord, electron microscopy shows enlarged bulbous neurites adjacent to motor neurons in the ventral horn (Fig. 5L), some of which show evidence of degeneration with collections of membranous debris and vacuolized mitochondria (Fig. 5M).

### Myelin deficits in R237H rat model of AxD

AxD is classically characterized as a leukodystrophy; nevertheless, little is known regarding how a primary disorder of astrocytes leads to deficits in white matter and myelination. R237H rats have noticeably smaller spinal cords compared to wild-type littermates, and area measures from cross sections of the cervical cord enlargement (C5) reflect this size difference (Fig. 6A). However, separate measures of white and gray matter show a disproportionate reduction in white matter (Fig. 6A). To assess further whether R237H rats have myelin deficits, we focused on spinal cord as a severely affected CNS region. Immunoblotting for myelin proteins shows both MBP (myelin basic protein) and CNP (a2’,3’-cyclic nucleotide 3’ phosphodiesterase) are modestly but significantly reduced in R237H rats at 8 weeks of age, while PLP1 (proteolipid protein) does not change (Fig. 6B). Electron microscopy shows thinner myelin sheaths and occasional degenerating axons with organelle accumulation, dark cytoplasm, or emptied ballooning myelin sheaths in the dorsal corticospinal tract (Fig. 6C). To specifically measure myelin thickness in this region, we calculated myelin/axon g-ratios and found significant differences across axons of all calibers (Fig. 6D). Although the distribution of myelinated axon size was similar in mutant and wild-type spinal cord, R237H rats demonstrated an increased number of small unmyelinated axons (Fig. 6D). Measures in optic nerve also show differences in myelin thickness, with no significant change in size distribution or unmyelinated axon number (Fig. 6E, fig. S8).

**Fig. 6.**
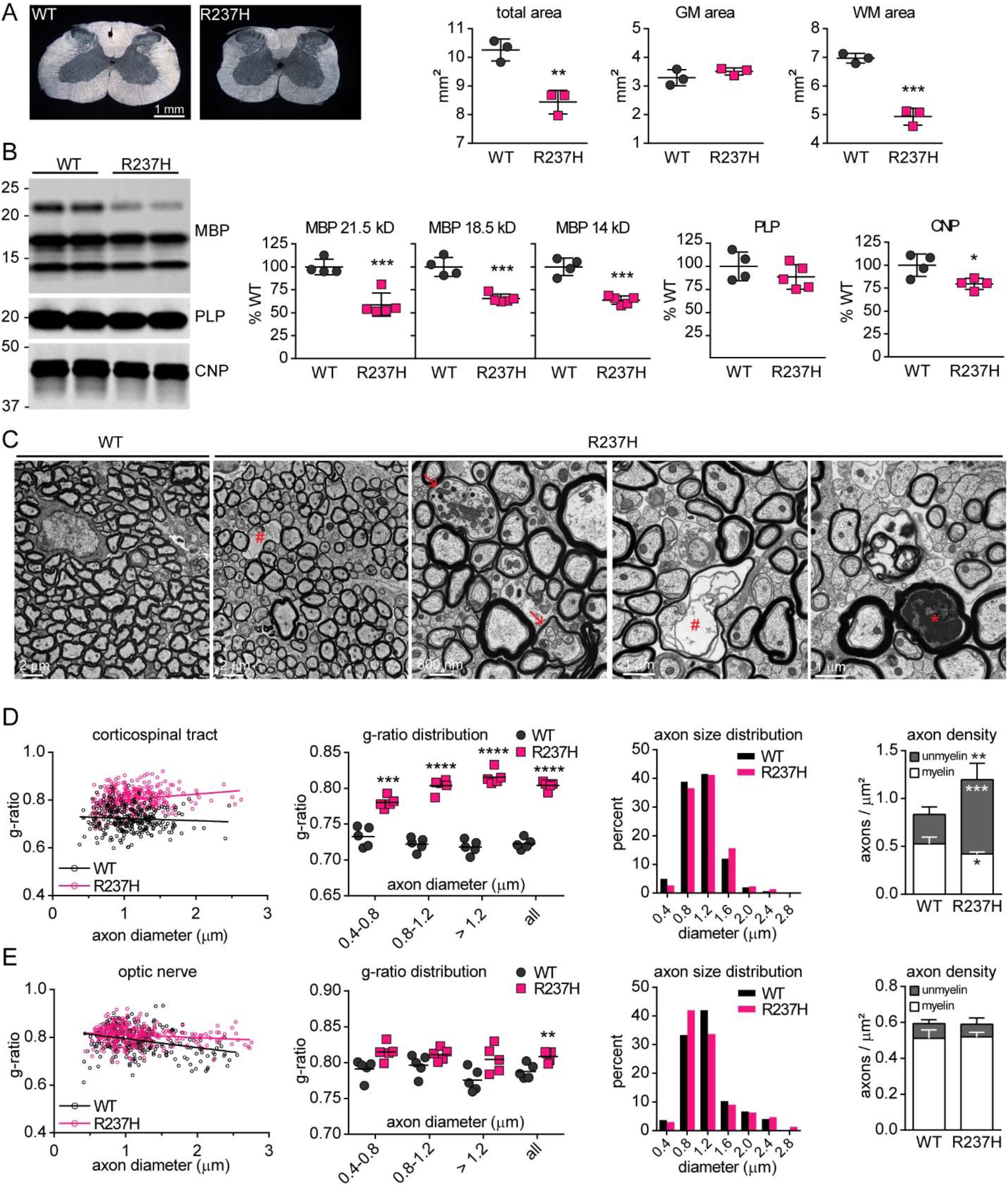
Myelin deficits in R237H rat model of AxD. (**A**) Area measures from cervical spinal cord cross sections (C5) show decreased size in R237H rats resulting from a general loss of white matter (WM) as opposed to gray (GM) at 8 weeks (males, N = 3). (**B**) Western analysis of cervical cord protein lysates shows a decrease in myelin proteins MBP and CNPase, but not PLP in R237H rats (two-tailed t-test, males and females at 8 weeks, N = 4-5). (**C**) Electron microscopy shows thinning myelin and degenerating axons with accumulating organelles (arrows), dark cytoplasm (asterisk), empty sheaths (hash symbol), and other peculiarities in the dorsal corticospinal tract. (**D**) Linear regression analysis of g-ratios shows significant difference in slopes between WT (−0.009997 ± 0.008478) and R237H (0.02370 ± 0.007269) cord (**p = 0.002595). Increased g-ratios confirm reduced myelin thickness across axons of all sizes (multiple t-tests, Holm-Sidak correction method, α = 5 %) with no change in myelinated axon size distribution (Kolmogorov-Smirnov test, p = 0.5197) but an increase in the number of unmyelinated axons (multiple t-tests as above). (**E**) Linear regression analysis of g-ratios in optic nerve shows thinner myelin sheaths (WT slope = −0.03716 ± 0.006892, R237H slope = −0.01209 ± 0.004445, **p = 0.001933) with no change in axon size distribution (Kolmogorov-Smirnov test, p = 0.1212) or the number of unmyelinated axons. N = 5 males per genotype, error bars = SD. For all tests: *p < 0.05, **p < 0.01, ***p < 0.001, ****p < 0.0001.

### GFAP suppression reverses AxD pathology

In comparison with mouse models, the AxD rat demonstrates more robust pathological phenotypes, and more importantly, clinically relevant motor deficits, which make it an ideal model to test therapeutic compounds. Two of the antisense oligonucleotides (ASO) designed to target mouse *Gfap* (*5*) cross react with the rat transcript. A dose response curve for the most potent ASO shows a reduction in rat *Gfap* transcript with increasing ASO concentrations (Fig. 7A). To determine the effects of incremental decreases in transcript on GFAP protein levels and composition, we fractionated lysates from cortex and cervical spinal cord from animals treated with increasing concentrations of ASO to quantify and compare soluble GFAP (monomers, dimers, tetramers), polymerized filaments, and Rosenthal fiber aggregates (Fig. 7B). The majority of GFAP exists in the filamentous cytoskeletal form (*19, 36, 37*). As expected, only mutant animals show significant amounts of GFAP in the Rosenthal fiber-enriched fraction. Although both the filamentous and aggregate forms are more resistant to biochemical solubilization, all three pools show a similar decrease with increasing doses of ASO, suggesting they are in equilibrium. GFAP levels in CSF and plasma also decrease with increasing dose of ASO (Fig. 7, C and D), again supporting potential utility as a biomarker.

**Fig. 7.**
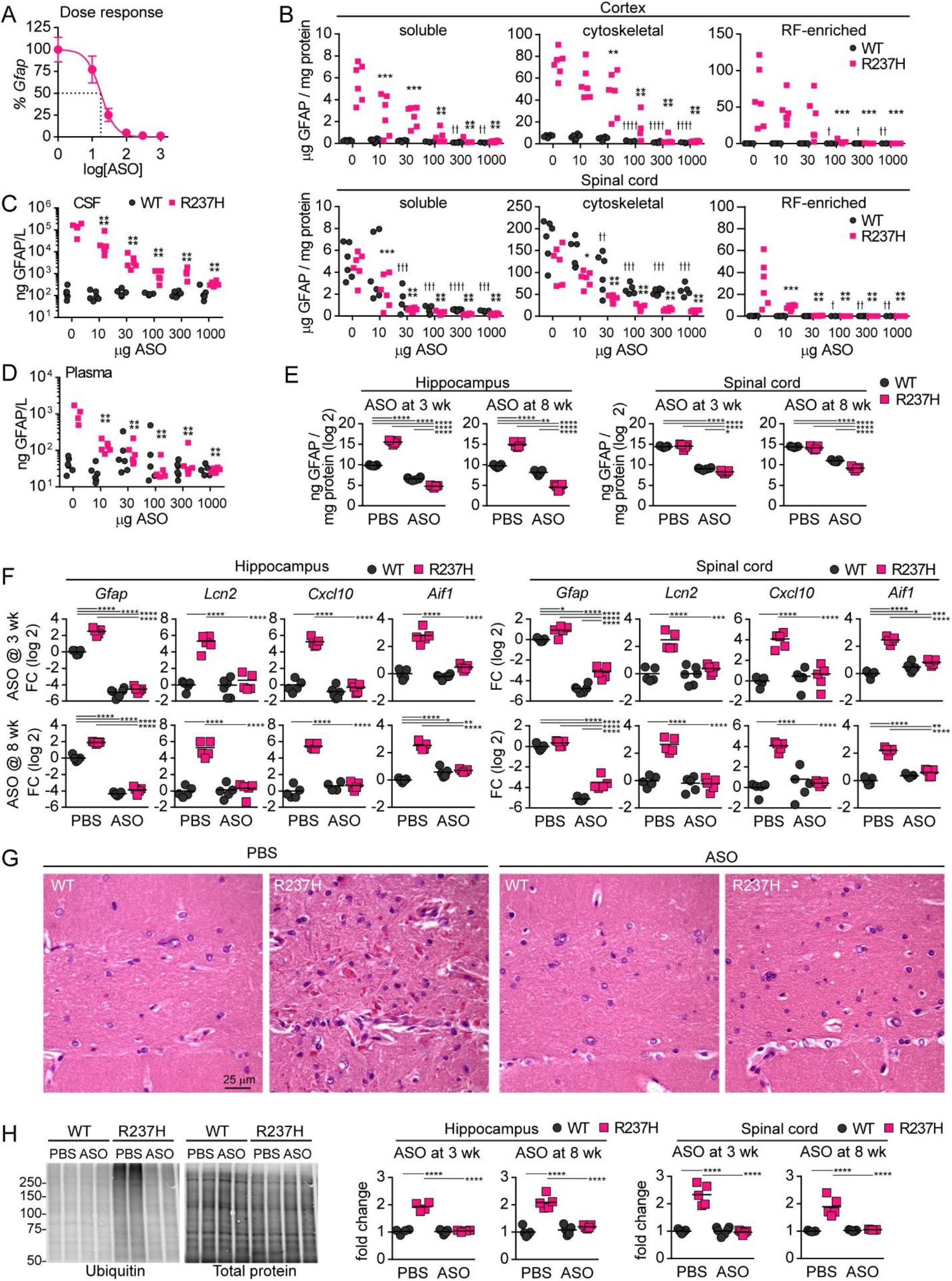
GFAP suppression reverses AxD pathology. (**A**) Dose response curve shows increasing concentrations of *Gfap*-targeted ASO ranging from 10 to 1000 µg lead to incremental reductions in *Gfap* transcript, as measured by qPCR in hippocampus from R237H rats, with a half-maximal effective concentration (EC50) of 18 µg (error = SEM). (**B**) Cortex and spinal cord protein fractions from the same cohort show dose dependent reductions in all forms of GFAP by ELISA. (**C** and **D**) Analysis of CSF (**C**) and plasma (**D**) show similar reductions in R237H rats and no reduction in WT rats (one-way ANOVA comparisons to vehicle treated group within genotypes, * indicates R237H, t indicates WT for Dunnett’s multiple comparisons test, N = 5-6). (**E**) Treatment at 3 and 8 weeks of age with 300 µg ASO for maximal effect suppresses total GFAP protein to levels below wild-type treated with PBS or ASO in R237H rats in hippocampus and spinal cord. (**F**) Transcript analysis (qPCR) shows reduction of *Gfap* and normalization of other markers of astro- and microgliosis in rats treated at 3 weeks or 8 weeks (**E** and **F**, two-way ANOVA of log2 transformed values with Tukey’s multiple comparisons, N = 5-6). (**G**) H&E shows reversal of Rosenthal fiber accumulation in ASO treated R237H rats (hippocampal fissure shown from rats treated at 8 weeks). (**H**) Western analysis demonstrates clearance of high molecular weight ubiquitinated proteins in hippocampus and spinal cord from R237H rats treated at either 3 weeks or 8 weeks. Western image shows representative hippocampal lysates from rats treated at 8 weeks, left panel is ubiquitin, right panel is total protein stain for normalization (two-way ANOVA with Tukey’s multiple comparisons, N = 5-6). For all post-test comparisons: ^*/✝^ p < 0.05, ^**/✝✝^ p < 0.01, ^***/✝✝✝^ p < 0.001, ^****/✝✝✝✝^ p < 0.0001.

To further test whether ASO treatment can prevent or reverse AxD pathology, we treated 2 cohorts of R237H rats and wild-type littermates with 300 µg ASO for maximal suppression (dose response in Fig. 7A): one at 3 weeks of age, immediately after weaning and before onset of obvious clinical phenotypes, and a second group at 8 weeks of age when animals are severely impaired (Fig. 1, F, H and I). Animals in both groups were tested for strength and coordination at 10 weeks post-treatment (13 and 18 weeks of age respectively), and tissues were collected 12 weeks post-treatment (15 and 20 weeks of age) to confirm GFAP suppression and assess cellular phenotypes. To confirm GFAP suppression, we quantified both transcript and total protein in hippocampus and spinal cord and found levels at or below the level of wild-type controls in both treatment groups (Fig. 7, E and F). Additional transcriptional markers associated with astrogliosis including lipocalin2 (*Lcn2*) and the small chemokine (*Cxcl10*) were normalized by ASO treatment. The marker for microglial activation Iba1 (*Aif1*) was also reduced in R237H treated animals compared to those receiving vehicle (Fig. 7F). Histological assessment confirmed clearance of Rosenthal fibers (Fig. 7G, hippocampus from rats treated at 8 weeks shown), and immunoblotting demonstrated clearance of high molecular weight ubiquitinated proteins regardless of treatment age (Fig. 7H). Furthermore, treatment of an additional cohort of animals at 3 weeks of age demonstrated sustained GFAP suppression out to 24 weeks post-treatment (Fig. S9).

### GFAP suppression rescues motor deficits and AxD phenotypes

We next asked whether the substantial correction in cellular and molecular pathology achieved by ASO administration yielded clinical benefit. After ASO treatment, animals were weighed weekly as a general measure of health and failure to thrive. R237H rats receiving ASO showed significant improvement in body weights. Those treated at 3 weeks showed no difference compared to wild-type littermates, and those treated at 8 weeks showed significant improvement compared to R237H rats receiving vehicle (Fig. 8A, Movie S3). R237H rats in both groups demonstrated increased grip strength and improved performance in the horizontal ladder test (Fig. 8, B and C), suggesting not only that deterioration could be prevented, but that functional deficits could be reversed by GFAP clearance. At the cellular level, the appearance of ramified microglia in ASO treated animals gave further evidence of a quieted neuroinflammatory response (Fig. 8, D to G), and in astrocytes, expression of proteins associated with normal astrocyte function such as KCNJ10 and AQP4 were normalized in rats of either age group (Fig. 8, H to O, spinal cord; rats treated at 8 weeks shown in D to O).

**Fig. 8.**
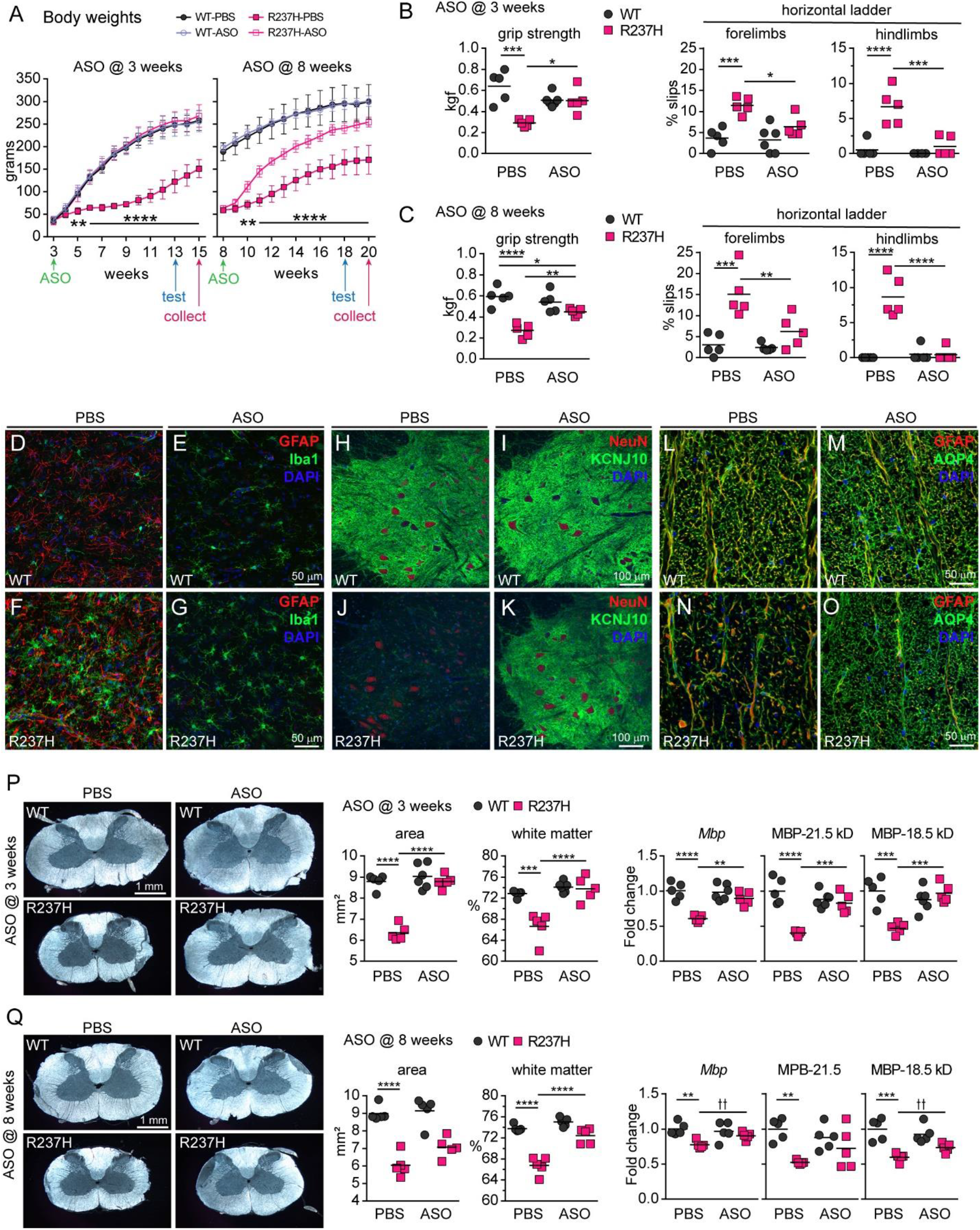
GFAP suppression rescues motor deficits and AxD phenotypes. (**A**) Body weights of rats treated at 3 or 8 weeks of age show significant differences between R237H rats receiving ASO compared to vehicle in both cohorts (repeated measures two-way ANOVA, Bonferroni’s multiple comparisons test, N = 5-6, error = SD). (**B** and **C**) Motor tests of forelimb grip strength (kgf = kilogram force) and coordination with paw placement on the horizontal ladder demonstrate prevention of motor deficits in animals treated at 3 weeks (**B**) and reversal of deficits in animals treated at 8 weeks (**C**) in all measures (two-way ANOVA, Tukey’s multiple comparisons test, N = 5-6). (**D** to **G**) Iba1 and GFAP immunofluorescent labeling (IF) shows GFAP suppression and restoration of ramified microglia in cervical spinal cord gray matter (ventral horn) from R237H rats treated at 8 weeks. (**H** to **O**) KCNJ10-IF (Kir4.1) demonstrates rescue of expression in ventral horn (**H** to **K**), and AQP4-IF shows increased labeling in white matter (**L** to **O**) in cervical cord from R237H rats treated at 8 weeks. (**P** and **Q**) Area measures of cervical cord cross sections show ASO treatment at 3 weeks prevents spinal cord white matter loss (**P**) and treatment at 8 weeks partially reverses white matter loss (**Q**). These improvements are reflected in MBP transcript and protein analysis in cervical cord for both cohorts (*p < 0.05, **p < 0.01, ***p < 0.001, ****p < 0.0001, two-way ANOVA, Tukey’s multiple comparisons tests; tt p < 0.01, multiple t-tests within genotypes, Holm-Sidak correction method, α = 5 %, N = 5-6).

With the rescue of both cellular and behavioral phenotypes, we wanted to know whether white matter changes in spinal cord could also be prevented or reversed (Fig. 8, P and Q). Area measures of spinal cord cross sections showed that treatment at 3 weeks of age prevented deterioration. While ASO treatment at 8 weeks of age did not show significant reversal of spinal cord size, the percentage of white matter increased. In addition, MBP transcript and protein levels increased in R237H rats treated with ASO at both time points.

## DISCUSSION

Alexander disease is a devasting neurological disorder, and GFAP suppression by targeted ASO therapy offers a real possibility for treatment. Our proof of concept studies using *Gfap* targeted ASOs in mouse models demonstrated reversal of AxD pathology, including protein accumulation and aggregation and the reactive stress response (*5*). However, AxD mice display relatively mild behavioral phenotypes, necessitating animal models with clinically relevant outcome measures in which to test functional improvement. The GFAP-R237H rat has a heavier burden of pathology with higher levels of GFAP, widespread Rosenthal fiber formation throughout the brain and spinal cord, myelin deficits and functional deficits including motor impairment, failure to thrive, and lethality.

In this report, the severe and progressive phenotype of the R237H rat allowed us to test two treatment strategies to either prevent or reverse behavioral deficits. Treatment of R237H rats with *Gfap*-targeted ASO at weaning cleared molecular and cellular pathology preventing onset of clinical phenotypes, and R237H rats treated at this age were physically indistinguishable from wild-type animals (Movie S3). More importantly, treatment of severely affected young adult rats not only cleared pathology, but partially reversed white matter deficits and motor impairment.

Recent studies have used ASO to specifically target neuronal and oligodendrocyte pathology in other degenerative disorders, including Huntington’s and Pelizaeus-Merzbacher disease respectively (*38, 39*). Here we demonstrate that targeting astrocyte specific *Gfap* rescues astrocyte function in AxD, allowing the recovery of other cell types in the CNS.

AxD is classified as a leukodystrophy, with frontal lobe white matter loss in patients with early onset and brainstem and cervical cord atrophy nearly universal in those with later onset (*1, 40–42*). To determine whether R237H rats have myelin deficits, we focused on cervical spinal cord as a relevant and discrete area for histological, molecular, and ultrastructural analysis. Reduced white matter volume, decreased myelin proteins, thin myelin sheaths and degenerating axons demonstrate clear deficits in R237H rat spinal cord, and thinner sheaths observed in optic nerve suggest these deficits may be more widespread. While astrocyte dysfunction may directly affect local myelination and spinal cord circuits, deterioration of other brain regions or descending spinal tracts may also contribute to cord atrophy and motor impairment. Future studies to assess CNS connectivity and descending versus ascending tracts in spinal cord, brainstem and other brain regions in the rat model could potentially reveal how astrocyte pathology affects long range relays, local circuits, or central pattern generators in AxD (*43–48*).

Astrocytes are vital in supporting white matter integrity by maintaining ion and neurotransmitter balance and through the secretion of a variety of myelin promoting and inhibiting factors, and other leukodystrophies including megalencephalic leukoencephalopathy with subcortical cysts (MLC) and vanishing white matter (VWM) disease are associated with astrocyte dysfunction (*49*). Altered expression and cellular distribution of SLC1A2, GJA1, AQP4, and KCNJ10, as observed in R237H rat astrocytes, could have profound effects on ion and neurotransmitter homeostasis and myelination. We have also reported that AxD associated mutations lead to increased brain stiffness (*22*), and tissue stiffness has been shown to regulate oligodendrocyte progenitor cell (OPC) proliferation and differentiation (*50*). In addition, recent studies with a human iPSC model of AxD show a potential connection between elevation of the astrocyte secreted glycoprotein CHI3L1 (chitinase 3 like 1, YKL-40) and reduced OPC proliferation (*51*). While it is still unclear how astrocyte dysfunction leads to white matter deficits in AxD and whether the leukodystrophy reflects hypo- or de-myelination, the rat model now offers the opportunity to study these effects in detail.

Early onset AxD is often associated with failure to thrive (*1*), and decreased body weight is a consistent feature among AxD rodent models (*7, 52*). While R237H rats achieve normal early postnatal milestones, they fail to gain weight after weaning and never reach the size of their wild-type littermates. Oropharyngeal dysphagia and frequent emesis are also common features in AxD that may contribute to body weight differences (*1, 53*) and remain to be investigated in the model. Although rodents cannot vomit (*54*), surrogate measures of nausea such as pica behavior (*55*) and treatment with antiemetics may inform whether brainstem pathology affects the chemoreceptor trigger zone in R237H rats. Intractable vomiting is also observed in neuromyelitis optica (NMO) spectrum disorders, where the autoimmune response against the high levels of APQ4 in circumventricular organs can lead to area postrema syndrome (*56*), suggesting astrocyte dysfunction may be a common denominator for frequent emesis in both AxD and NMO. Several studies have also tied astrocyte function to hypothalamic regulation of food intake and energy balance (*57–60*), and a detailed analysis of brainstem and hypothalamus will be essential in determining causes for failure to thrive and lethality in the model.

*Gfap*-targeted ASO treatment leads to long lasting suppression of GFAP transcript and protein to below wild-type levels in both mouse and rat models of AxD. We have found that ASO treatment can not only prevent, but also reverse, many aspects of disease, and that functional improvement is possible even when treatment is delayed until the animals are already severely impaired. Although the ASOs used in this study were selected to target rodent *Gfap*, efforts to move human *GFAP* targeted ASO into clinical trials are underway. While we are able to achieve nearly complete knockdown of GFAP in the models, the same level of suppression may not be possible in human, and future efforts will focus on determining the level of reduction necessary to relieve relevant phenotypes in the R237H rat.

## MATERIALS AND METHODS

### Study design

Our goal for this study was to develop a preclinical model of AxD with relevant pathological and behavioral phenotypes to facilitate studies of pathophysiology and testing of novel therapeutics. The targeted mutation in the rat GFAP gene was based on the severity and frequency of the R239H mutation in human disease. Rats were used for their increased potential for functional phenotypes compared to the mouse, and the wealth of literature on behavioral research. Generation of the R237H rat was contracted through Applied StemCell, Inc. (Milpitas, CA), and first-generation offspring were bred and analyzed at the University of Wisconsin – Madison. All animal studies were approved by the College of Letters and Sciences and Vice Chancellor’s Office for Research and Graduate Education Animal Care and Use Committee.

Experimental approach was based on the following goals: 1) to confirm the targeted mutation and to breed transgenic animals to wild-type rats to remove potential off-target mutations; 2) to confirm the presence of hallmark AxD pathology including Rosenthal fibers and the astrocyte stress response; 3) to identify cell- and non-cell-autonomous effects of GFAP mutation; 4) to identify behavioral phenotypes relevant to AxD; 5) to test *Gfap*-targeted ASOs to validate the model and as a promising therapeutic to prevent or rescue behavioral phenotypes.

Animal numbers for each experiment were determined by our previous experience with mouse models of AxD. Both sexes were used for experiments, and in some cases, females were used preferentially due to their smaller size for long-term co-housing. ASO treatment ages and endpoints were determined prospectively based on the progression of the rat phenotype and time course studies in the mouse. A small number of R237H rats were removed from studies due to death or euthanasia of moribund animals. Rats were randomly assigned sequential numerical identifiers before weaning and genotyping, and treatment assignments were alternated within sex and genotype groups in the order of the identification number. No more than two animals per group (sex, genotype, treatment) were taken from a single litter. Outliers were only excluded if the result reflected technical failure (e.g. low normalization signal indicating sample loss).

Laboratory staff performing behavioral assays or measurements were blinded to genotype and treatment, and those performing molecular and histological analyses were blinded until after data collection when possible. Given the severity of the pathology and physical phenotype of the R237H rat, blinding was not always possible.

### Statistical analysis

Technical replicates were included for all molecular analyses (qPCR, ELISA), and averaged per animal. All data points are shown with the mean when possible. Statistics are based on animal numbers per group and do not include technical replicates. Typically, two tailed t-tests were used for comparisons of two groups, one-way ANOVA with Tukey’s (all groups compared) or Dunnett’s (groups compared to control) multiple comparisons for comparisons of multiple groups under the same conditions, multiple t-tests for comparisons of different measures between two groups, two-way ANOVA with Tukey’s multiple comparisons for tests with two conditions (e.g. genotype, treatment), and two-way repeated measures ANOVA with Sidak’s multiple comparisons to compare two groups over time. Tests used are indicated in the figure legends.

GraphPad Prism v6 was used for statistical analysis and graphing.

## Supplementary Materials

Materials and Methods

Fig. S1. Complementary DNA sequence of *Gfap* modifications in CRISPR/Cas9 targeted rat lines.

Fig. S2. Reduced white fat in R237H rats.

Fig. S3. STAT3 activation in astrocytes of R237H rats.

Fig. S4. Widespread reactive gliosis in R237H rat brain.

Fig. S5. Rosenthal fibers in R237H rats.

Fig. S6. Early caspase-3 activation in R237H astrocytes and TUNEL positive cells in adult rats.

Fig. S7. Astrocyte membrane proteins in total protein lysates.

Fig. S8. Myelin deficits in optic nerve.

Fig. S9. GFAP suppression at 24 weeks post-ASO-treatment.

Table S1. Antibodies used in the study.

Table S2. Primers used in the study.

Move S1. Wild-type rat (female) on horizontal ladder at 8 weeks of age.

Move S2. R237H rat (female) on horizontal ladder at 8 weeks of age.

Move S3. Alexander model rats (male) treated with ASO at P21.

## Acknowledgments

We would like to thank Denice Springman, Rebecca Pulvermacher, Jacob Loeffelholz, and Alder Levin for technical support.

## Author contributions

T.L.H, B.P., M.B.F., M.-D.P, R.F.B., and A.M designed the study and participated in preparation of the manuscript, T.L.H and A.F.M. performed initial RNA and protein analysis and immunohistochemistry to validate the model, N.-H.L. and M.-D.P. prepared protein for all other immunoblot analysis of astrocyte proteins, K.L.D performed myelin protein analysis, other immunoblotting in ASO treated animals, and immunohistochemistry. S.H. prepared protein lysates and fractions and performed ELISA assays for GFAP and rat IgG. T.L.H., S.H., and A.M analyzed EM micrographs for g-ratio determination.

## Data and materials availability

All data associated with this study are present in the paper or the Supplementary Materials. The mutant rats will be deposited with the Rat Resource and Research Center at the University of Missouri in Columbia, MO.

