## Supplemental materials for "Antisense therapy in a new rat model of Alexander disease reverses GFAP pathology, white matter deficits, and motor impairment"

### SUPPLEMENTARY MATERIALS AND METHODS

#### R237H rat production and housing

A point mutation was engineered at the endogenous rat *Gfap* locus using CRISPR-Cas9 technology to produce the equivalent of the severe Arg239His mutation found in many human AxD patients (Arg237His in the rat GFAP sequence, due to minor species differences in the N-terminal head domain). A microinjection cocktail consisting of rat GFAP RNA (5'-TCGCACTCAGTACGAGGCAGTGG-3'), single-strand donor oligonucleotide, and Cas9 mRNA (each at 50 ng/µl) was injected into the cytoplasm of single-cell fertilized embryos from Sprague-Dawley rats. Founders carrying the desired mutation were identified by PCR analysis of DNA isolated from tail biopsies, using a pair of primers that flank exon 4 (Table S1), which are predicted to yield a 769 bp product after amplification. After the F1 generation, animals were backcrossed to the CD®(SD)IGS-001 strain (code 001, hereafter abbreviated CD-001), with one generation derived from a wild-type male sire to transfer the wild-type Y chromosome. Animals are genotyped by Transnetyx (Cordova, TN) or a sequence specific TaqMan probe (Thermo Fisher Scientific, Applied Biosystems). Data described in this report are from animals representing at least 4 generations of backcrossing to CD-001, except in initial experiments to confirm the targeted mutation and protein expression (Fig. 1). Percent survival was assessed retrospectively from animals scheduled for experiments at 12 weeks or later as well as those held as breeders.

#### Transcript analysis of CRISPR/Cas9 targeted *Gfap*

To confirm that the mutant *Gfap* allele was expressed, RNA was isolated from brain and converted to cDNA as described below for quantitative PCR. Primers spanning exons 2-5 (Table

S1) were used to PCR amplify the targeted exon 4 mutation for initial Sanger sequence analysis. PCR products were purified with the Wizard SV Gel and PCR Clean-Up System (Promega) and diluted to 10 ng/100 bp in a 20 µl BigDye Terminator v3.1 Cycle Sequencing reaction (Thermo Fisher Scientific) with 25 ng primer (Table S1), and sequenced on a 3730xl Genetic Analyzer (Thermo Fisher Scientific, Applied Biosystems) at the University of Wisconsin – Madison Biotechnology Center. Primers spanning 2,475 bp including the ATG start site to the 3'UTR (Table S1) were used to amplify the full coding sequence with Platinum Taq DNA Polymerase High Fidelity and cloned into the with the TOPO TA Cloning Kit (pCR™2.1-TOPO vector, Thermo Fisher Scientific, Invitrogen), according to the manufacturer's protocol, to verify sequence integrity and full expression of the targeted allele.

**RNA isolation and quantitation**

Tissues were collected and frozen immediately with dry ice or liquid nitrogen and subsequently homogenized in Trizol reagent (Thermo Fisher Scientific, Invitrogen) for RNA isolation per the manufacturer's protocol. For quantitative PCR, 1 µg RNA was transcribed as cDNA using 1 µm random hexamers, 2 µm poly-(dT)15 primers and 500 nm dNTP in a 20 µl reaction with Superscript III (Thermo Fisher Scientific, Invitrogen protocol). Reverse transcriptase reactions were diluted and incorporated into quantitative PCR at a ratio of 16 nl cDNA per µl reaction using SYBR Green Master Mix with 150 nM primers (Table S1) on an ABI ViiA7 Real-Time PCR System (Thermo Fisher Scientific, Applied Biosystems Inc). Standards generated from purified amplicons were used to determine relative concentrations and values were normalized to *Rn18s*. *Mbp* transcripts were quantified with a TaqMan probe (Thermo Fisher Scientific, ABI assay ID Rn01399619_m1) using Fast Advanced Master Mix and normalized to *Ppia* (Rn00690933_m1) using the ΔΔCt method.

**Preparation of CNS lysates and immunoblotting for astrocyte-related proteins**

Total brain lysates (Fig. 1C, 2A to B) were prepared by homogenizing half brains bisected at the sagittal midline in SDS lysis buffer (50 mM, Tris-HCl (pH 7.4), 5 mM EDTA, 2% (w/v) SDS, 1 mM Pefabloc SC (Sigma-Aldrich) and Complete$^{TM}$ Protease Inhibitor Cocktail (Sigma-Aldrich, Roche) at 50 mg/ml using a rotor-stator homogenizer. After homogenization, protein concentration was determined by bicinchoninic acid assay using the BCA Protein Assay kit (Thermo Fisher Scientific) with bovine serum albumin (BSA) as a standard. To fractionate brain lysates for further analyses (Fig. 3, A to B), rat brain regions and cervical spinal cord were homogenized with 10 volume per weight (100 mg/ml) of radioimmunoprecipitation (RIPA) buffer (20 mM Tris, pH 7.4, 150 mM NaCl, 1% (v/v) Triton X-100, 5 mM EDTA, 0.5% (w/v) sodium deoxycholate, and 0.1% (w/v) SDS) containing cocktails of protease inhibitors (10 µM ALLN, 2 µg/ml leupeptin, 5 µg/ml aprotinin, and 2 mM PMSF, all from Sigma-Aldrich) and phosphatase inhibitors (5 mM sodium fluoride, 1 mM sodium vanadate, 1 mM sodium pyrophosphate, and 1 mM β-glycerol phosphate, all from Sigma-Aldrich). Samples were centrifuged at 17,000 × g for 20 min at 4°C and the supernatant was designated as the RIPA-soluble fraction. Pellets were homogenized in urea buffer (6M urea, 5 mM EDTA, and 20 mM Tris, pH 7.4) and centrifuged at 17,000 × g for 20 min at 4°C. The supernatant was designated as the urea-soluble cytoskeletal fraction. The insoluble pellets were solubilized by resuspension in 1% SDS buffered by 10 mM Tris-HCl, pH 8 and 5 mM EDTA and boiled for 5 min. Membrane fractions, as well as nuclear and cytosolic fractions (Fig. 2D and 4A) were prepared by

Membrane and Nuclear/Cytosol Fractionation kits (BioVision, Milpitas, CA), according to the manufacturer's instructions. After protein concentration determination by BCA assay (Thermo Fisher Scientific), protein samples were mixed with appropriate volumes of sample buffer (25 mM Tris-HCl, pH 6.8, 10% (v/v) glycerol, 1% (w/v) SDS, and 5% (v/v) β-mercaptoethanol) prior to analysis by immunoblotting.

Proteins (10-15 μg) were analyzed by immunoblotting using 12% (w/v) SDS-PAGE and nitrocellulose membranes (Pall Life Sciences). After electrophoretic transfer of protein, the membranes were blocked with 3% (w/v) BSA in Tris-buffered saline with Tween (TBST: 20 mM Tris-HCl, pH 7.4 and 150 mM NaCl, containing 0.1% (v/v) Tween 20) at room temperature for 1 hour. After being washed with TBST three times, membranes were incubated with primary antibody (Table S2) at 4°C overnight, followed by incubation with horseradish peroxidase-conjugated anti-mouse, anti-rabbit or anti-rat secondary antibodies (Table S2). The blots were developed with Enhanced Chemiluminescence substrate (Western Lightning, Perkin Elmer Life Sciences), and signals were digitized using a LAS 4000 Imaging Analyzer (GE Healthcare). Signals from nonsaturated exposures of immunoblots were quantified using the ImageQuant software (Ver. 7.0, GE Healthcare).

**Protein preparation and myelin protein analysis**

Frozen tissues were processed in ice-cold RIPA buffer (Thermo Fisher Scientific, Pierce) with 1mM PefablocSC (Sigma-Aldrich), and Complete Protease Inhibitors (Roche) at 50 mg/ml with a rotor-stator homogenizer. Samples were centrifuged at 17,000 g for 20 min at 4°C, and supernatants collected for protein quantitation with the BCA assay (Thermo Fisher Scientific, Pierce). Protein concentrations were adjusted for equivalent gel loading and unsaturated signal as follows, MBP = 1 µg, PLP = 0.1 µg, CNPase = 5 µg per lane (Fig. 6B). Samples were combined with 2x Laemmli buffer (BioRad), heated to 95°C for 10 min (MBP, CNP) or 37°C for 30 min (PLP to prevent aggregation), loaded on 12 % (CNP) or 4-20 % gradient (MBP, PLP) Criterion TGX gels (BioRad), electrophoresed, and transferred to Immobilon-FL PVDF membrane (Millipore Sigma IPFL00010). Transferred protein was visualized with REVERT 700 Total Protein Stain (LI-COR), imaged and quantified with a LI-COR Odyssey system. Immunoblots were subsequently blocked in Sea Block (Thermo Fisher Scientific, Pierce 37527) for 1-2 hours before incubating in primary antibody diluted in TBS with 0.05% Tween20 (Thermo Fisher Scientific, Pierce 28360) overnight at 4°C. Membranes were washed in TBS/Tween20 before incubation with secondary antibodies at room temperature for 2 hours, washing (TBS/Tween20 followed by PBS) and Odyssey imaging. Antibody details for MBP, PLP, CNPase, and GAPDH are listed in Table S2. PLP was normalized to GAPDH due to the low amount of total protein analyzed while MBP and CNP were normalized to total protein signal.

**Western analysis for ASO treated animals**

Total protein was prepared from snap frozen tissues by homogenizing with a bead mixer mill (GenoGrinder, SPEX SamplePrep) at 50 mg/ml 2% SDS in 50 mM Tris-HCl, 5 mM EDTA pH 7.4, 1mM PefablocSC (Sigma-Aldrich), and Complete Protease Inhibitors (Roche). Samples were boiled for 20 min and protein quantified using the BCA assay (Thermo Fisher Scientific, Pierce). To quantify high molecular weight ubiquitinated proteins (Fig. 7H), 15 µg total protein were loaded per lane on 10 % Criterion TGX gels, processed as indicated above, and probed with antibodies for ubiquitin (Table S2). For MBP detection (Fig. 8, P and Q), 0.8 µg total protein were loaded on a 4-20% gradient gel, transferred to Immobilon-FL PVDF membrane with

(*61*)buffer for high PI proteins, and processed as indicated above for MBP and GAPDH protein (Table S2). Ubiquitin signal was normalized to total protein and MBP was normalized to GAPDH.

**GFAP protein quantification by ELISA**

GFAP was quantified from either total protein extracted with 2% SDS as described above or from biochemical fractions separating soluble, filamentous, and aggregate forms of GFAP (*19*) by a sandwich enzyme linked immunosorbent assay (ELISA) as described previously (*5*). Briefly, microtiter plates are coated with a cocktail of monoclonal GFAP antibodies diluted in PBS (SMI-26, see Table S2 for antibody details) and blocked with BLOTTO (5% milk in PBS) before adding protein samples and standards diluted in PBS with 1% BSA and 0.05% Tween 20 (Tw20). Plates are incubated at room temperature for 2 hours and washed with PBS/0.05% Tw20 before adding rabbit anti-GFAP detection antibody diluted in BLOTTO (Dako Z0334) and incubating overnight at 4°C. Plates are washed with PBS/0.05% Tw20 followed by incubation with secondary HRP conjugated goat anti-IgG diluted in BLOTTO at room temperature for 2 hours. Plates are washed with PBS/0.05%Tw20 followed by PBS before adding SuperSignal ELISA Femto Substrate (Thermo Fisher Scientific, Pierce) and reading with a GloRunner luminometer (Turner Biosystems).

For GFAP protein fractions, tissues were homogenized at 50 mg tissue/ml in 0.5% Triton-X-100/ 20mM Tris-HCl/ 2mM EDTA pH 7.4 (TEE) with 1mM PefablocSC and Complete Protease Inhibitor Cocktail, using a bead mixer mill (GenoGrinder), and centrifuged at 17,000 g for 20 min. at 4°C. The Triton-X-100 soluble supernatant was collected, and the insoluble pellet was resuspended in an equivalent volume of 6M urea in TEE and centrifuged at

3000 g for 10 minutes at 4°C. The supernatant was collected as the urea soluble filamentous fraction, and the RF-enriched pellet was resuspended in 2% SDS in TEE and boiled for 30 minutes. Triton-X-100 soluble lysates were assayed at 0.3 µg/ml, urea soluble fractions at 0.005 (cord) – 0.03 (cortex) µg/ml, and SDS soluble fractions at 0.05 µg/ml. To quantify GFAP from total protein lysates, 2% SDS homogenates were diluted to 0.1 to 0.2 µg/ml.

For body fluid analysis, blood was collected transcardially into EDTA coated tubes (BD Microtainer tubes, 365974), centrifuged at 1,000 g for 20 min at 4’C, and GFAP quantified from plasma diluted 1:1 with 1% BSA in PBS/ 0.05% Tw20. CSF collected through the cisterna magna was diluted in a range of 1:4 to 1:16 in 1% BSA in PBS/ 0.05% Tw20 for GFAP quantitation as described previously (*20*).

A portion of samples from ASO treated animals were below the limit of detection at the protein concentrations used in the ELISA. For statistical analysis, GFAP concentrations for these samples were set to the lowest quantity from the standard curve and values extrapolated per mg protein in the assay.

**Immunohistochemistry and imaging**

For immunofluorescent labeling of tissues, animals were deeply anesthetized with isoflurane before transcardial perfusion of PBS followed by 4% paraformaldehyde for fixation. Brain and spinal cord were collected and post-fixed overnight before cryoprotecting in graded concentrations of sucrose (10-30%) and sectioning on a sliding microtome at 40 µm. Sections were stored at -20°C in 25 % glycerol/ 25 % ethylene glycol/ 0.1 M phosphate buffer as cryoprotectant. For labeling, tissue sections were rinsed with PBS, blocked and permeabilized with 5 % normal goat or donkey serum, 0.5% Triton-X-100 in PBS, and incubated in primary antibody (Table S2) diluted in 1% BSA, 0.3 % TX100 in PBS 72 hrs at 4°C. Sections were washed in PBS with 0.05% TX100 before incubating in secondary antibody diluted in 1% BSA, 0.3 % TX100 in PBS at 4°C overnight. Finally, sections were washed in PBS/0.05% TX100 and mounted with ProLong Gold mounting media with DAPI (Thermo Fisher Scientific, Invitrogen). For phophoY705-STAT3 immunofluorescence, sections were further permeabilized with 100% methanol for 10 min at -20°C prior to adding block. For GFAP immunohistochemistry, sections were treated with 0.15 % $H_2O_2$ to block endogenous peroxidases before blocking and incubating with mouse anti-GFAP (Table S2, Sigma G6171) as above, and VECTASTAIN Elite ABC-HRP Kit (peroxidase, mouse IgG, PK-6102) was used in combination with Vector SG substrate for chromogen labeling according to the manufacturer's protocol (Vector Laboratories). Fluorescence images were collected with a Nikon A1R-HD confocal microscope system with equivalent settings for comparisons between genotype or treatment groups. Brightfield images were taken with a SPOT camera (Diagnostic Instruments) on a Nikon Microphot.

**TUNEL labelling**

Apoptotic cell death was visualized in 6 μm sections from paraffin embedded tissue using TUNEL labelling according to the manufacturer's instructions (TdT FragEL DNA fragmentation kit, Calbiochem), with an additional avidin-biotin-peroxidase amplification step and the substrate diaminobenzidine (Vector Laboratories). Astrocytes were identified by co-labeling for GFAP using the mouse monoclonal anti-GFAP antibody N206/8 (Neuromab) at a dilution of 1:500, a secondary anti-mouse alkaline phosphatase coupled antibody (SouthernBiotech), and the alkaline phosphatase substrate Vector Red (Vector Laboratories).

**Spinal cord imaging and area measures**

For spinal cord collections, segments were identified by vertebral number and spinal nerve location and cervical cord segment C5 isolated for sectioning after perfusion with fixative (see above). Frozen sections collected at 30 µm with a sliding microtome were mounted on slides with Prolong Gold (Thermo Fisher Scientific, Invitrogen) and imaged by phase contrast (Ph4) with a SPOT camera (Diagnostic Instruments) on a Nikon Microphot. Image J was used to trace white and gray matter to calculate percentages and total area.

**Electron microscopy and g-ratio analysis**

Animals were perfused with PBS followed by 2 % paraformaldehyde/2.5 % glutaraldehyde in 0.1 M phosphate buffered saline. Spinal cords were removed after dorsal laminectomy and C5 identified by spinal nerve location. Optic nerves were collected after cutting at the base of the eye and then removing the brain and cutting at the optic chiasm. Nerves were trimmed at the midpoint between the eye and chiasm for cross sections. Tissues remained in fixative until osmication and embedding at the University of Wisconsin Medical School Electron Microscopy Facility, and ultrathin sections were viewed with a Philips CM120 scanning transmission electron microscope (UW-Madison, WI).

To quantify myelin sheath thickness and g-ratios, 3 representative images (5600x) were taken from grids of the dorsal corticospinal tract at C5 or optic nerve from 5 male rats at 8 weeks for each genotype. From each image, 20 axons were selected using a random number generator for X-Y image coordinates for a total of 60 axons per animal. Myelin thickness was determined by tracing the outer and inner edge of the myelin sheath in Fiji/ImageJ and calculating the diameter from the area assuming a circular shape, $d = 2 \times \mathrm{SQRT}(\mathrm{area}/\pi)$. G-ratios were then calculated as the inner/outer myelin diameter. To quantify myelinated and unmyelinated axons, a 12 $\mu m^2$ grid was superimposed on the same images and the Cell Counter Plugin was used in Fiji/ImageJ to count axons in every other grid frame (9 per image) using the principles of stereology to avoid over counting (i.e. for axons that cross grid lines, only those that crossed the top and right lines were counted while those crossing the left and bottom lines were not).

**Postnatal developmental milestone assessment**

Developmental milestones were assessed to track the growth and development of the newborn rats. Milestones were evaluated on post-natal days (PND) 4, 6, 8, 10, and 12 for 78 rats from 15 litters. Measurements were taken of body temperature, weight, body length, and tail length. Other milestones include the righting reflex, circle traverse, negative geotaxis, and cliff avoidance. Righting reflex is tested by laying the rat pups on their back and timing how long it takes them to right themselves onto all four paws, with a maximum of 60 seconds. For the circle traverse, the pup was placed into the center of a circle 12.5 centimeters in diameter. Pups had a maximum of 30 seconds to get at least half of their body out of the circle. To measure negative geotaxis, pups were placed facing down approximately ¼ of the way from the bottom of a 34 cm long by 23 cm wide wire mesh screen that was slanted downwards at an angle of 45 degrees. Pups had a maximum of 60 seconds to turn around and climb up the wire mesh screen. Cliff avoidance was measured by placing the front two paws over the edge of a table. Pups had a maximum of 60 seconds to turn away from the edge and avoid falling. Values from animals of each genotype were averaged per litter as a unit for statistics.

**Behavior tests in juvenile and adult rats**

The open field task was used to measure locomotor activity, anxiety, and exploration levels of the rats. Illumination of the test room was set at approximately 25-30 lux. Rats were tested twice in open field, once on PND 21, and again on PND 52. For this test, rats were placed in the test arena (42 x 42 x 30 cm) for 30 minutes. Integra software was used to track the movements of the rats. Data were then exported and analyzed for multiple aspects of activity, including horizontal activity, vertical activity, total activity, and time spent in the center.

Grip strength was measured with a meter from Columbus Instruments equipped with a horizontal grid pull-bar. Animals were positioned above the meter and allowed to grasp the grid with their forepaws before pulling them horizontally by the tail away from the meter until they released the grid. Peak force was recorded for 5 trials with approximately 10 minutes between trials. The lowest and highest measures were excluded and the remaining 3 values averaged for comparisons.

Performance on the accelerating rotarod (Med Associates) was assessed over 4 trials consisting of the rod accelerating from 5 to 40 RPM over a 5-minute period. The duration of time spent on the rod before falling was recorded for each trial with a minimum of 20 minutes between each trial.

The horizontal ladder consists of two Plexiglas walls (1 m x 20 cm) spaced 10 cm apart with rungs at the bottom (2 cm spacing) elevated 30 cm from the benchtop for animals to traverse (*62, 63*). Rats were placed at the beginning of the apparatus and allowed to move freely across while paw placement was video recorded. A food reward was placed at the end of the ladder for animals tested at 8 weeks, and for older animals (15-20 weeks), the home cage was placed at the end. Steps and paw slips were recorded for each limb after the first full stride with slips consisting of complete misses or slips off the rungs. Animals were tested over 3 trials approximately 20 minutes apart.

**CNS water content**

To measure water content in brain and spinal cord, 8-week-old male rats were euthanized by $CO_2$ asphyxiation, the brain and cervical cord were removed and microdissected on ice to isolate brainstem, cerebellum, hippocampus, cortex, and corpus callosum, which were then snap frozen in liquid nitrogen. CNS regions were subsequently weighed on glass slides (wet weight), desiccated at 100°C for 72 hours, and weighed again (dry weight) to determine water content. Percentages were calculated as the wet-dry weight difference divided by the wet weight.

**Blood brain barrier permeability**

To assess the blood brain barrier, we measured rat IgG by ELISA (Immunology Consultants Laboratory, E-25G) as a blood protein not normally found in brain parenchyma and an indicator of barrier breakdown (*64*). Female WT and R237H rats at 10 weeks of age (N = 8) were anesthetized with isoflurane and transcardially perfused with saline until blood was cleared completely from the animal (0.5 ml saline per gram). Hippocampus and cervical spinal cord were collected immediately, frozen on dry ice, and stored at -80 °C prior to analysis. Tissues were homogenized with a bead mixer mill (GenoGrinder, SPEX SamplePrep) at 100 mg/ml RIPA buffer (Thermo Fisher Scientific, Pierce) with Complete Protease Inhibitor Cocktail (Sigma-Aldrich) and 5 mM EDTA. Samples were centrifuged at 14,000 g for 20 minutes at 4 °C and the supernatant collected for analysis. Protein was quantified using the BCA assay (Thermo Fisher Scientific, Pierce) and samples diluted to a concentration of 50 μg/ml with diluent provided by the Rat IgG ELISA Kit. The assay was performed according to the manufacturer's protocol and absorbance measured at 450 nM with a VersaMax microplate reader (Molecular Devices). Concentration of IgG was determined using the standard curve with a sigmoidal 4 parameter logistic curve and then normalized to the total protein concentration of each sample.

**ASO treatment**

Animals were treated with ASO by intracerebroventricular (ICV) injection as described previously for mice (*5*) with minor modifications of stereotaxic coordinates and injection volume. Briefly, rats are deeply anesthetized with isoflurane and placed into a stereotaxic apparatus to inject 30 µl ASO in phosphate buffered saline or PBS alone. For rats at P21, the injection coordinates were +0.2 mm anterior of bregma -1.5 mm lateral of the midline and -3 mm ventral from the surface of the skull. For adult wild-type rats (8-10 weeks), the coordinates were -1 mm posterior, -1.5 mm lateral, and -3.7 mm ventral. For adult R237H rats, the coordinates used were the same as those for P21 rats. In dose response experiments, WT and R237H adult rats (10 weeks, males and females, N = 6) were treated with 0, 10, 30, 100, 300, and 1000 µg ASO in a 30 µl volume, and tissues collected 8 weeks post-injection for transcript and protein analysis to assess suppression. For behavioral experiments, female rats (N=5) were treated at P21 or P56 for maximal suppression with 300 µg ASO, motor tests performed 10 weeks post-treatment and tissues collected 12 weeks post-treatment. Animals were euthanized by $CO_2$ asphyxiation and brain and cervical cord removed. Brains were bisected at the midline with the ipsilateral side immersion fixed in 4% paraformaldehyde and the contralateral side partitioned into brain regions and frozen for transcript and protein analysis. Spinal cord was similarly divided with C4-C6 placed in fixative and C1-3 frozen.

### SUPPLEMENTARY FIGURES

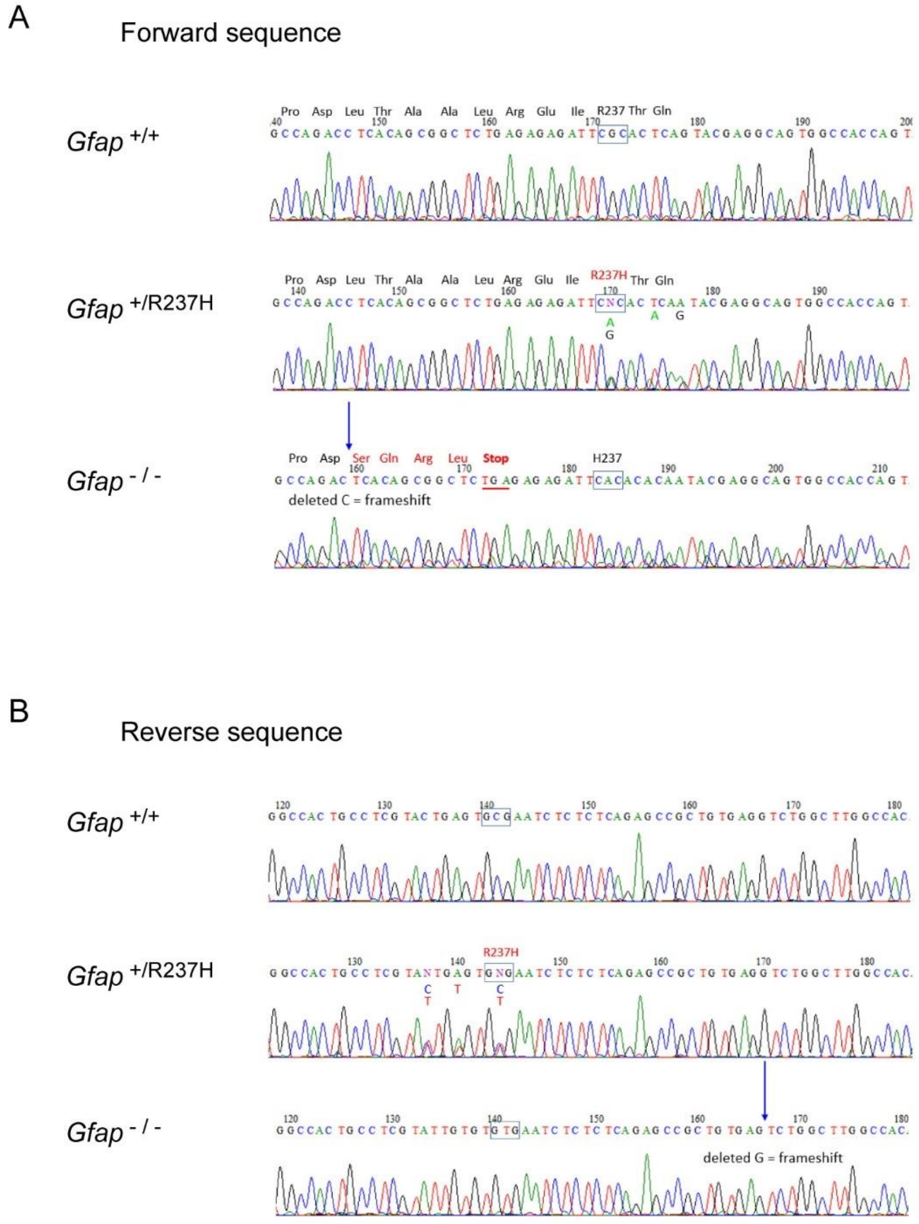


**Fig. S1. Complementary DNA sequence of *Gfap* modifications in CRISPR/Cas9 targeted rat lines.** Sequence analysis of PCR amplified cDNA from two different founder lines compared to wild-type (*Gfap*$^{+/+}$) rats shows the correct sequence leading up to the Arg237 codon and targeted His modification in *Gfap*$^{+/R237H}$ rats compared to the single nucleotide deletion and subsequent frame shift in addition to the targeted R237H mutation in the second knockout line (*Gfap*$^{-/-}$). Both forward (**A**) and reverse (**B**) sequences are shown. Note that the two silent mutations following the R237H codon were introduced to block re-cutting and improve accuracy of homology-directed repair.

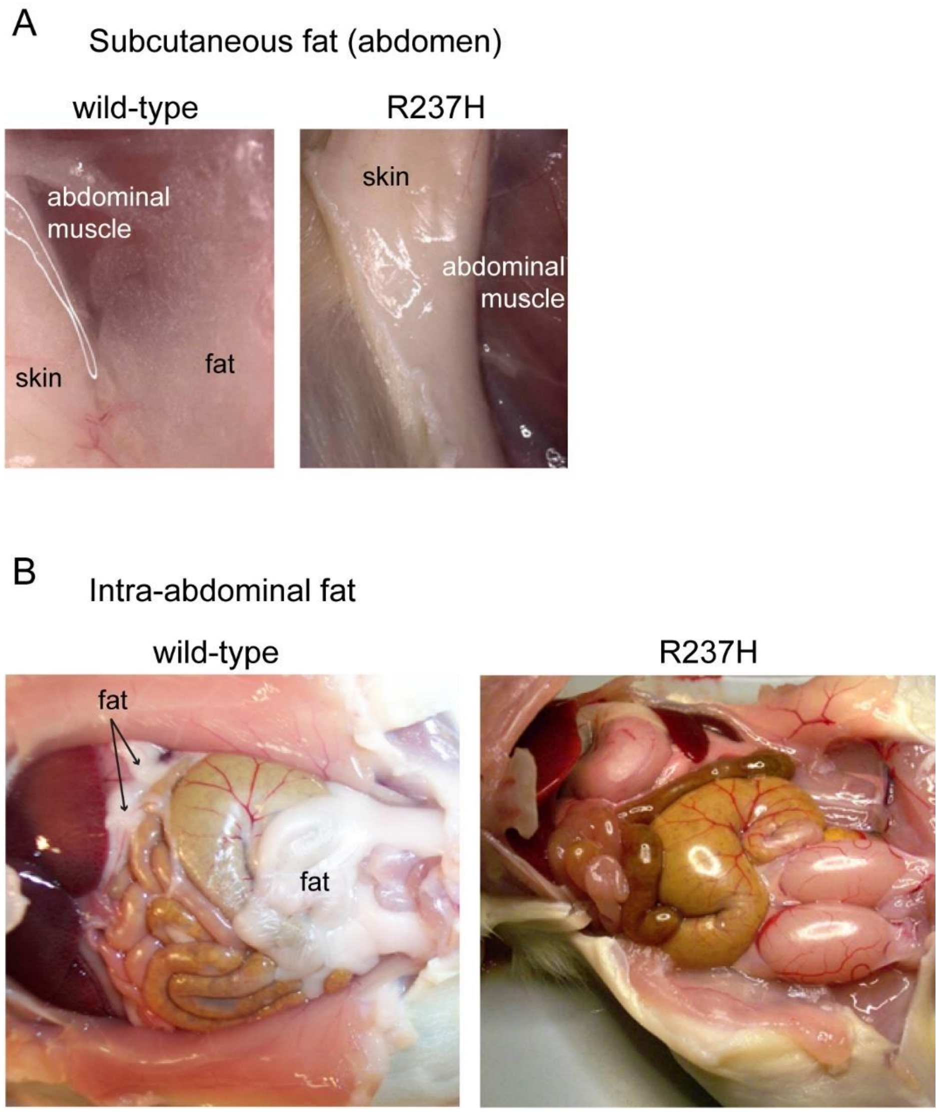


**Fig. S2. Reduced white fat in R237H rats.** At 8 weeks of age, R237H rats exhibit virtually no white fat. (**A**) shows subcutaneous fat overlying the abdominal musculature in a WT rat and its absence in R237H rats. (**B**) shows the accumulation of intra-abdominal fat in WT compared to R237H animals.

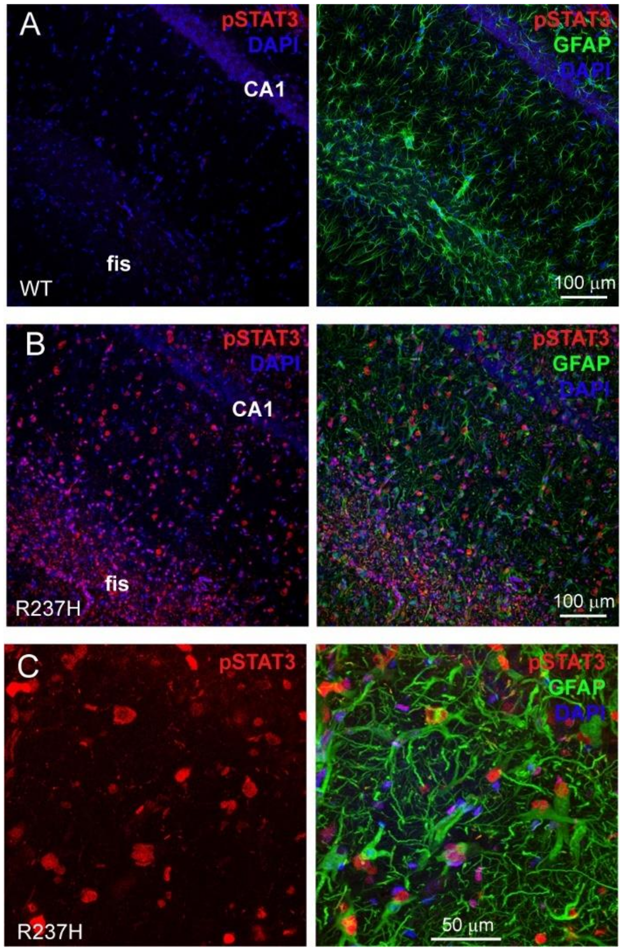


**Fig. S3. STAT3 activation in astrocytes of R237H rats**. (**A** to **C**) Immunolabeling shows nuclear localization of phospho-Y705-STAT3 in GFAP labeled astrocytes and potentially other cell types in hippocampus from R237H rats (**B** and **C**) compared to WT (**A**). Hippocampal fissure (fis) and CA1 regions indicated in **A** and **B**, while **C** shows higher magnification of CA1.

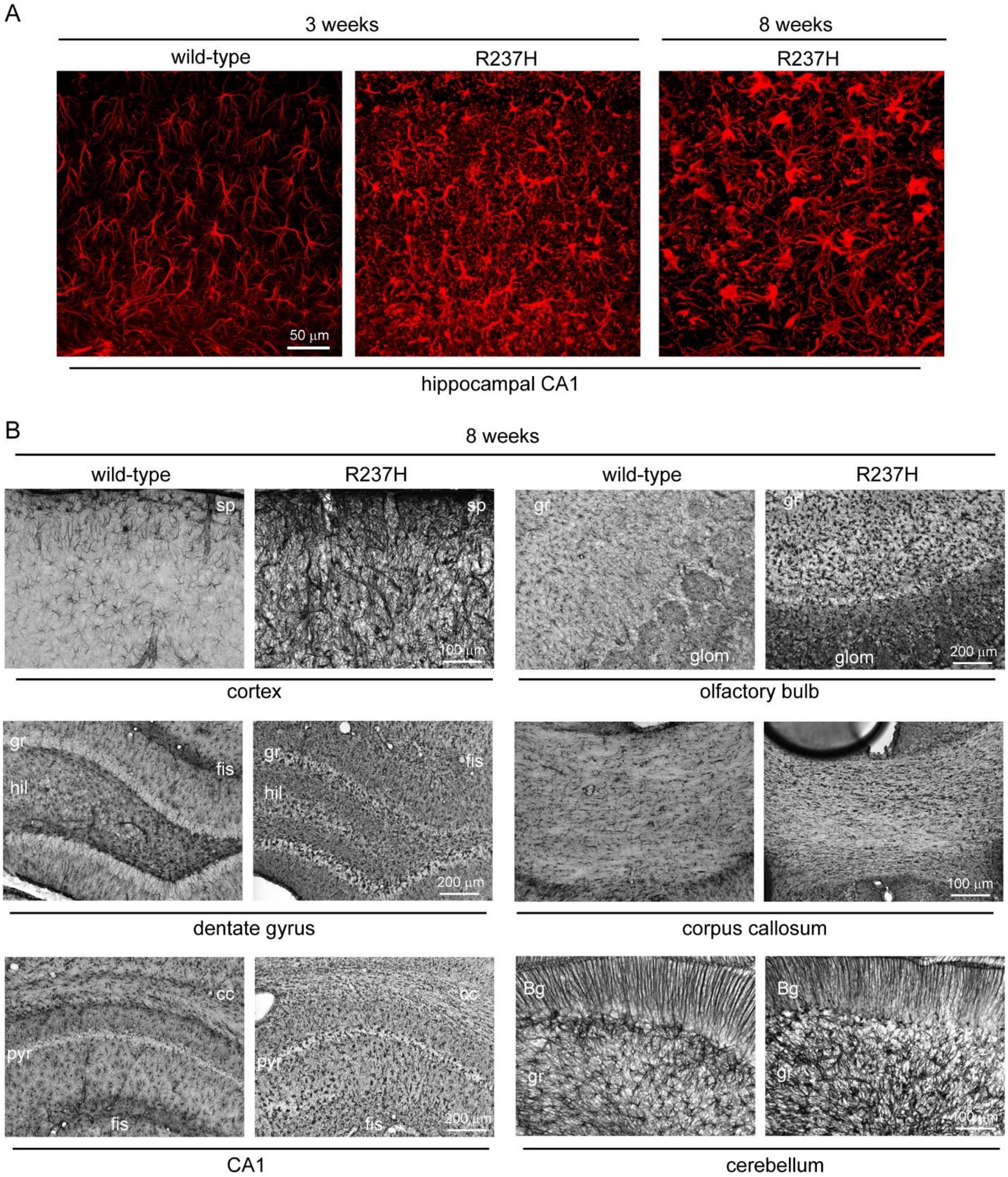


**Fig. S4. Widespread reactive gliosis in R237H rat brain**. (**A**) GFAP immunolabeling at 3 weeks of age shows hypertrophied astrocytes with elevated GFAP in R237H rats, and at 8 weeks, astrocytes are globular and dysmorphic (hippocampal CA1 shown). (**B**) Reactive astrocytes with elevated GFAP are apparent throughout the CNS with generalized gliosis in many regions. While astrocytes in the cerebellar granule cell layer and white matter also appear reactive, Bergmann glia maintain their radial morphology. sp = subpial, gr = granule cell layer, glom = glomerular layer, hil = hilus, fis = hippocampal fissure, pyr = pyramidal cell layer, Bg = Bergmann glia.

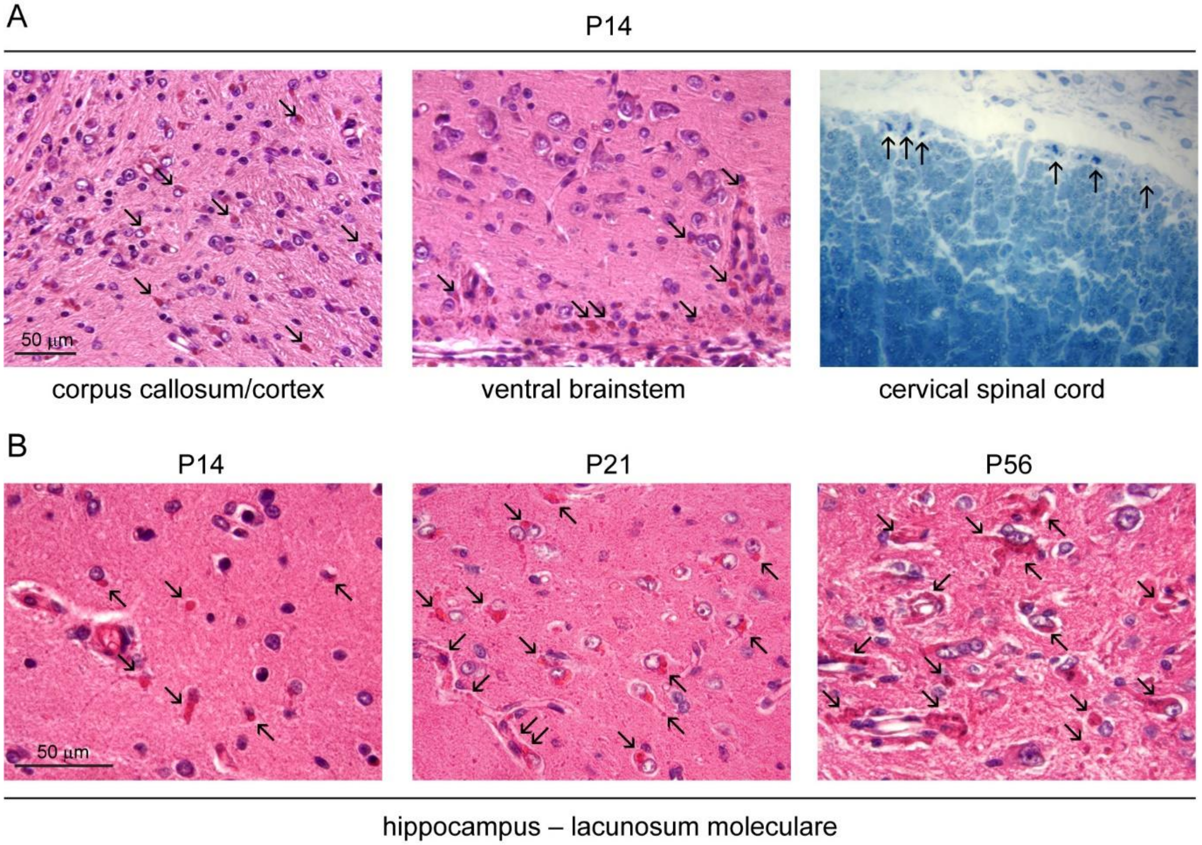


**Fig. S5. Rosenthal fibers in R237H rats.** (**A**) Rosenthal fibers are apparent as early as P14 as demonstrated by eosinophilic inclusions in H&E stained cortex, corpus callosum and brainstem as well as toluidine blue-stained spinal cord (arrows). (**B**) RFs become more prevalent as the animals age as shown in the stratum lacunosum-moleculare of the hippocampus (H&E).

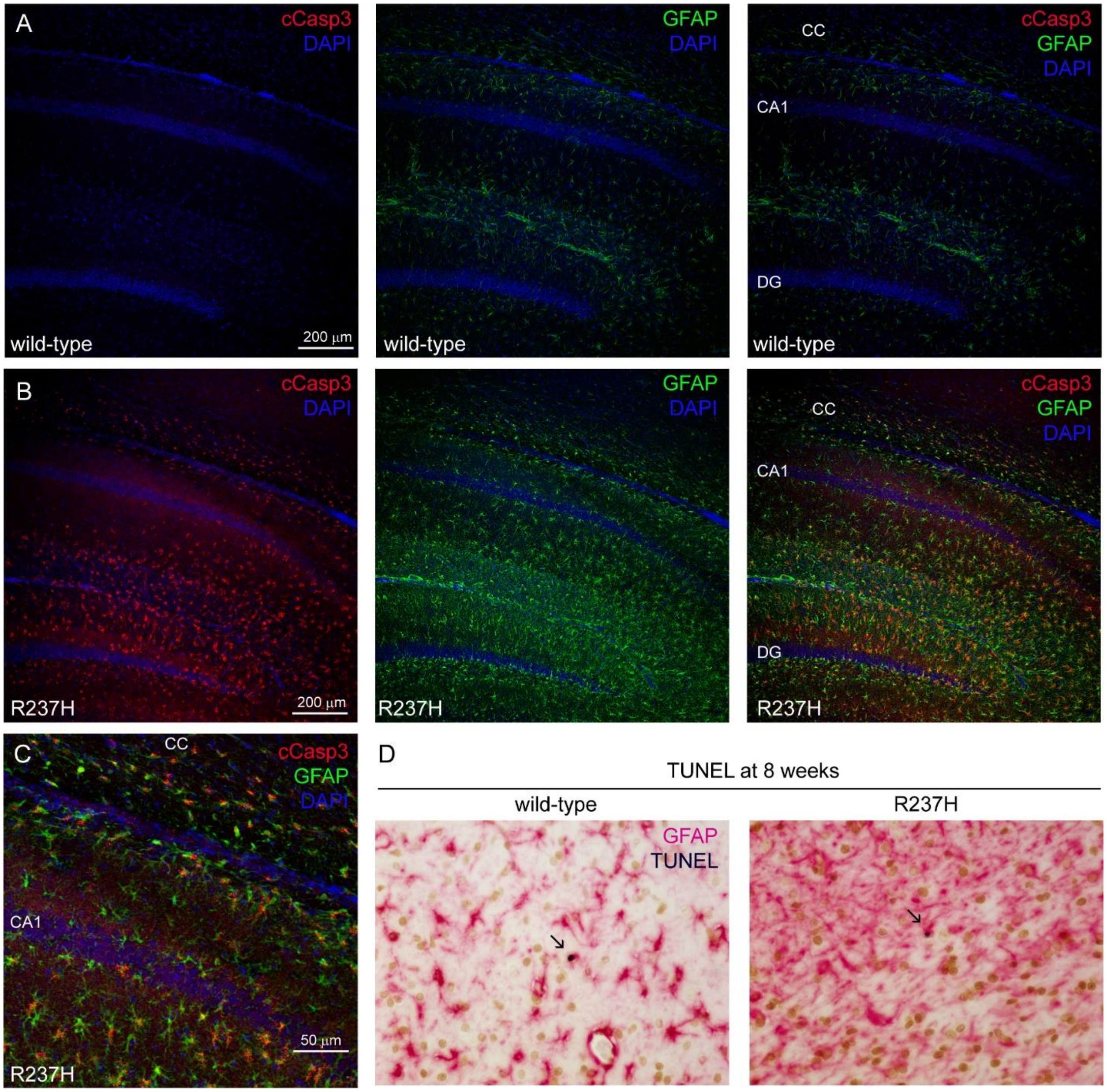


**Fig. S6. Early caspase-3 activation in R237H astrocytes and TUNEL positive cells in adult rats.** (**A** to **C**) Cleaved caspase-3 is prominent in R237H rats (**B**) at 3 weeks of age compared to WT animals (**A**, hippocampus shown, DG = dentate gyrus, CC = corpus callosum). (**C**) Higher magnification shows caspase-3 activation in GFAP labeled astrocytes. (**D**) TUNEL positive cells are apparent in wild-type rats but more frequent in R237H rats, although not all TUNEL labeled cells can be identified as astrocytes with GFAP co-labeling.

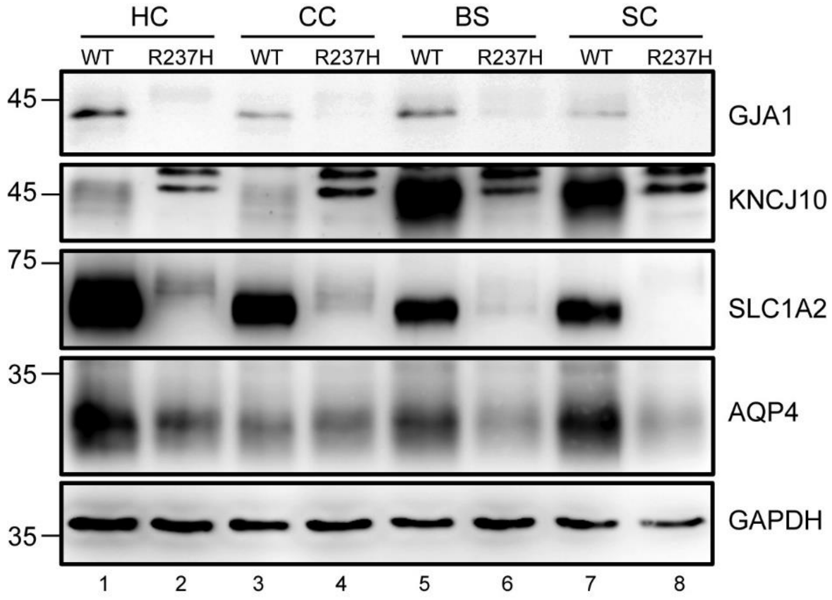


**Fig. S7. Astrocyte membrane proteins in total protein lysates**. Immunoblotting analysis for membrane proteins using SDS total protein lysates from different brain regions and spinal cord shows a similar pattern of expression as membrane fractionation (Fig. 4A). Higher molecular weight bands for KCNJ10 (Kir4.1) suggest this protein may be affected by post-translational modifications in the R237H rat. Although additional experiments would be necessary to confirm the signal is specific, the shift in localization observed with immunofluorescence, particularly in hippocampus (Fig. 4J and K), support the suggestion of aberrant protein modifications.

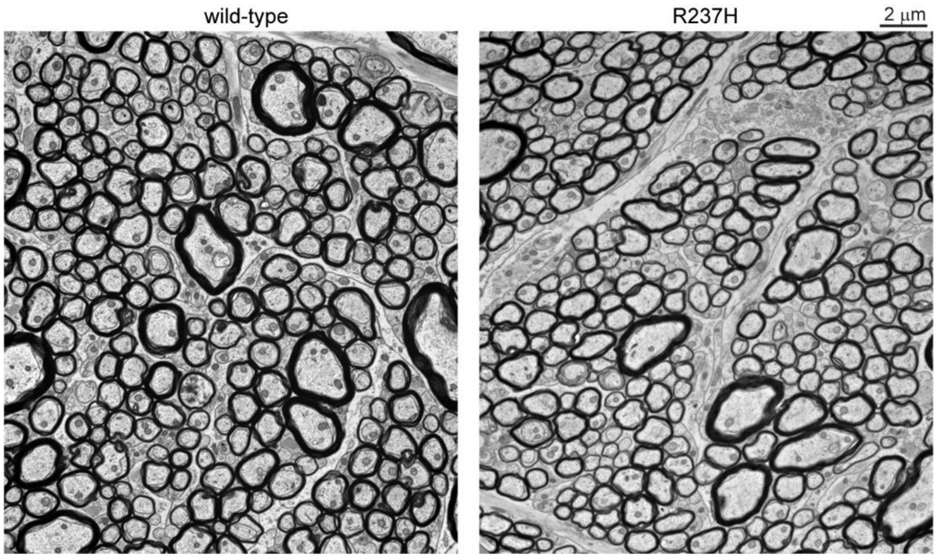


**Fig. S8. Myelin deficits in optic nerve.** Electron microscopy shows subtle differences in myelin thickness in R237H rats compared to WT. Representative images used for g-ratio analysis (Fig. 6E) are shown.

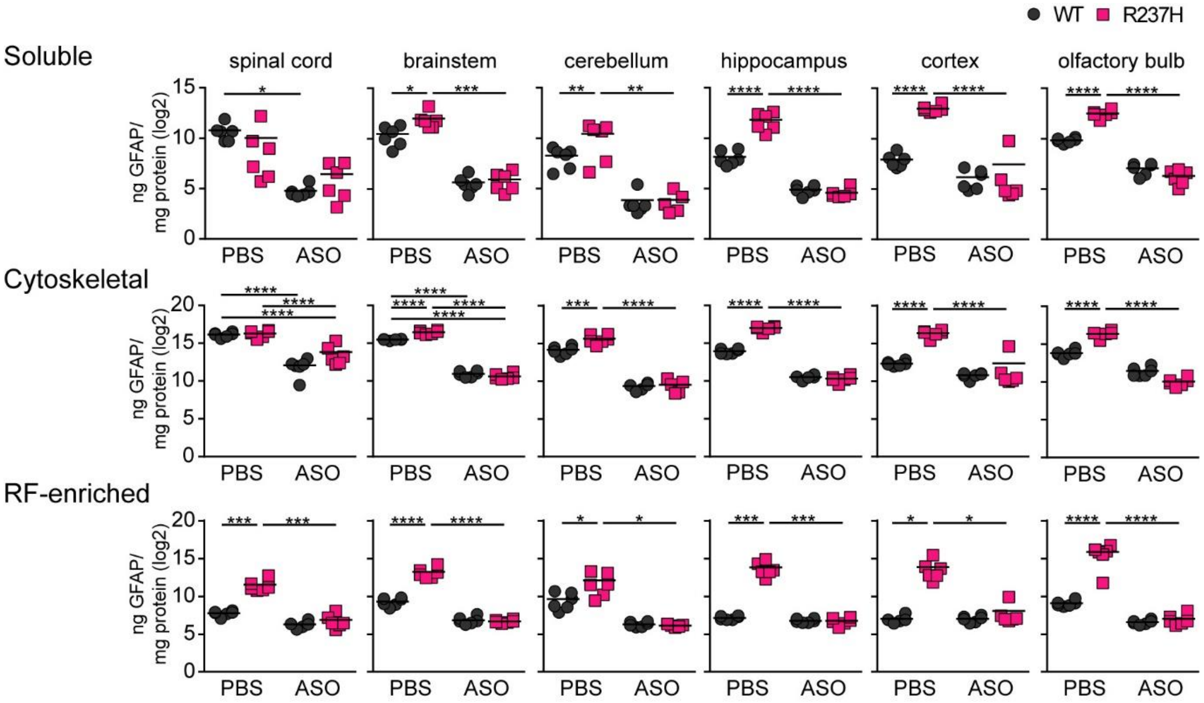


**Fig. S9. All forms of GFAP remain suppressed at 24 weeks post-ASO-treatment.** Biochemical fractions of GFAP including soluble, cytoskeletal (filamentous) and RF-enriched forms, show sustained reductions in all CNS regions analyzed in ASO treated R237H rats 24 weeks after treatment at 3 weeks of age (*$p < 0.05$, **$p < 0.01$, ***$p < 0.001$, ****$p < 0.0001$, two-way ANOVA, Tukey's multiple comparisons, N = 6 per group, 3 males, 3 females), with the sole exception of soluble GFAP in R237H spinal cord.

Table S1. Antibodies used in this study

| Experiment | Antigen | Host species | mono/poly | Source | Catalog No | RRID | Dilution |
|---|---|---|---|---|---|---|---|
| ELISA | GFAP | mouse | mono SMI 26 | BioLegend | 837602 | AB_2565380 | 1:1000 |
| ELISA | GFAP | rabbit | poly | Dako/Agilent | Z0334 | AB_10013382 | 1:5000 |
| IF | AQP4 | rabbit | poly | Millipore Sigma | AB3594 | AB_91530 | 1:500 |
| IF | Cleaved caspase-3 | rabbit | mono 5A1E | Cell Signaling Technology | 9664 | AB_2070042 | 1:400 |
| IF | CRYAB | mouse | mono 1B6.1-3G4 | Enzo Life Sciences | SPA-222 | AB_1659585 | 1:100 |
| IF | DCX | goat | poly | Santa Cruz Biotech | sc-8066 | AB_2088494 | 1:500 |
| IF | GFAP | mouse | mono GA5 | Millipore Sigma | G6171 | AB_1840893 | 1:1000 |
| IF | GFAP | rabbit | poly | Dako/Agilent | Z0334 | AB_10013382 | 1:1000 |
| IF | GFAP | rat | mono 2.2B10 | Virginia Lee (Univ Penn) | | | 1:200 |
| IF | GJA1 (CX43) | mouse | mono RN26 | Millipore Sigma | MAB3068 | AB_2110187 | 1:400 |
| IF | Iba1 | rabbit | poly | Wako | 019-19741 | AB_839504 | 1:500 |
| IF | KCNJ10 (Kir4.1) | rabbit | poly | Alomone Labs | APC-035 | AB_2040120 | 1:2000 |
| IF | NeuN | mouse | mono A60 | Millipore Sigma | MAB377 | AB_2298772 | 1:500 |
| IF | NG2 | rabbit | poly | Millipore Sigma | AB5320 | AB_91789 | 1:200 |
| IF | OLIG2 | mouse | mono 211F1.1 | Millipore Sigma | MABN50 | AB_10807410 | 1:100 |
| IF | SOX9 | goat | poly | R&D Systems | AF3075 | AB_2194160 | 1:200 |
| IF | SQSTM1 (p62) | mouse | mono 2C11 | Abnova | H00008878-M01 | AB_437085 | 1:100 |
| IF | STAT3-pY705 | rabbit | mono D3A7 | Cell Signaling Technology | 9145 | AB_2491009 | 1:200 |
| IF | Ubiquitin | rabbit | poly | Leica Biosystems/Nova Castra | NCL-UBIQ | AB_564048 | 1:500 |
| Western | AQP4 | rabbit | poly | Millipore | MAB3594 | AB_91530 | 1:5000 |
| Western | CCND2 | rabbit | mono D52F9 | Cell Signaling Technology | 3741S | AB_2070685 | 1:5000 |
| Western | CNPase | mouse | mono SMI 91 | BioLegend | 836404 | AB_2566639 | 1:500 |
| Western | CRYAB | mouse | mono 1B6.1-3G4 | Enzo Life Science | SPA-222 | AB_1659585 | 1:5000 |
| Western | GAPDH | mouse | mono 6C5 | Fitzgerald | 10R-G109A | AB_1285808 | 1:10,000 |
| Western | GAPDH | rabbit | poly | Abcam | Ab9485 | AB_307275 | 1:2500 |
| Western | GAPDH | mouse | mono 1D4 | Novus | NB300-221 | AB_350535 | 1:5000 |
| Western | GFAP | mouse | mono GA5 | Millipore Sigma | G6171 | AB_1840893 | 1:1000 |
| Western | GFAP | rabbit | poly | Dako/Agilent | Z0334 | AB_10013382 | 1:5000 |
| Western | GFAP C-term | mouse | mono N206A/8 | NeuroMab | 73-240 | AB_10672298 | 1:5000 |
| Western | GFAP N-term | mouse | mono SMI-21 | Covance | SMI-21R-100 | AB_509978 | 1:5000 |
| Western | GJA1 (CX43) | mouse | mono 4E6.2 | Millipore | MAB3067 | AB_94663 | 1:5000 |
| Western | HSPB1 | rabbit | poly | Enzo Life Science | SPA-801 | AB_1193479 | 1:5000 |
| Western | KCNJ10 (Kir4.1) | rabbit | poly | Alomone | APC-035 | AB_2040120 | 1:5000 |
| Western | Lamin A/C | mouse | mono 4C11 | Cell Signaling Technology | 4777S | AB_10545756 | 1:5000 |
| Western | MBP | goat | poly | Santa Cruz Biotech | sc-13914 | AB_648798 | 1:1000 |
| Western | NFkB-p65 | rabbit | mono D14E12 | Cell Signaling Technology | 8242S | AB_10859369 | 1:5000 |
| Western | NFkB-p65-pS536 | rabbit | mono 93H1 | Cell Signaling Technology | 4025S | AB_10827881 | 1:5000 |
| Western | PLP1 | rabbit | poly | Abcam | ab28486 | AB_776593 | 1:1000 |
| Western | SLC1A2 (GLT1) | guinea pig | poly | Millipore | MAB1783 | AB_90949 | 1:5000 |
| Western | SQSTM1 (p62) | mouse | mono 2C11 | Abnova | H00008878-M01 | AB_437085 | 1:5000 |
| Western | STAT3 | mouse | mono 124H6 | Cell Signaling Technology | 9139S | AB_331757 | 1:5000 |
| Western | STAT3-pY705 | rabbit | mono D3A7 | Cell Signaling Technology | 9145S | AB_2799407 | 1:5000 |
| Western | Ubiquitin | mouse | mono Ubi-1 | Millipore Sigma Chemicon | MAB1510 | AB_2180556 | 1:10,000 |
| Western | Ubiquitin | mouse | mono P4D1 | Cell Signaling Technology | 3936S | AB_331292 | 1:5000 |
| Western | VIM | rabbit | mono D21H3 | Cell Signaling Technology | 5741S | AB_10695459 | 1:5000 |

Table S1. Antibodies used in this study (continued)

| Experiment | 2nd antigen | Host | Conjugate | Source | Catalog No | Identifier | Dilution |
|---|---|---|---|---|---|---|---|
| ELISA | rabbit-IgG | goat | HRP | Millipore Sigma | A6154 | AB_258284 | 1:10,000 |
| IF | mouse-IgG | goat | Alexa Fluor 568 | Thermo-Invitrogen | A11031 | AB_144696 | 1:500 |
| IF | rabbit-IgG | goat | Alexa Fluor 488 | Thermo-Invitrogen | A11034 | AB_2576217 | 1:500 |
| IF | rat-IgG | goat | Alexa Fluor 647 | Thermo-Invitrogen | A21247 | AB_141778 | 1:500 |
| IF | goat-IgG | donkey | Alexa Fluor 488 | Thermo-Invitrogen | A11055 | AB_2534102 | 1:500 |
| IF | goat-IgG | donkey | Alexa Fluor 546 | Thermo-Invitrogen | A11056 | AB_2534103 | 1:500 |
| IF | mouse-IgG | donkey | Alexa Fluor 546 | Thermo-Invitrogen | A10036 | AB_2534012 | 1:500 |
| IF | mouse-IgG | donkey | Alexa Fluor 647 | Thermo-Invitrogen | A31571 | AB_162542 | 1:500 |
| IF | rabbit-IgG | donkey | CF-568 | VWR-Biotium | 20098 | AB_10557118 | 1:500 |
| IF | rabbit-IgG | donkey | Alexa Fluor 488 | Thermo-Invitrogen | A21206 | AB_2535792 | 1:500 |
| Western | mouse-IgG | goat | Alexa Fluor 680 | Thermo-Invitrogen | A21057 | AB_2535723 | 1:10,000 |
| Western | mouse-IgG | goat | DyLight 800 | Thermo Scientific | SA5-10176 | AB_2556756 | 1:10,000 |
| Western | rabbit-IgG | goat | Alexa Fluor 680 | Thermo-Invitrogen | A21109 | AB_2535758 | 1:10,000 |
| Western | rabbit-IgG | goat | DyLight 800 | Thermo Scientific | 35571 | AB_614947 | 1:10,000 |
| Western | goat-IgG | donkey | IRDye 800CW | LI-COR | 925-32214 | AB_2687553 | 1:10,000 |
| Western | mouse-IgG | Goat | HRP | Jackson ImmunoResearch Lab | 115-035-003 | AB_10015289 | 1:5,000 |
| Western | rabbit-IgG | Goat | HRP | Jackson ImmunoResearch Lab | 111-035-003 | AB_2313567 | 1:5,000 |
| Western | guinea pig-IgG | Goat | HRP | Jackson ImmunoResearch Lab | 106-035-003 | AB_2337402 | 1:5,000 |

TableS2. Primers used in this study

| Gene | Primer 1 | Primer 2 | Purpose |
|---|---|---|---|
| *Gfap* (intron 3 to 4) | AGTAACATCTGCCTCATTTCCGT | TGGCCTCATCTGGACTAAAGAAC | DNA amplifcation, sequencing |
| *Gfap* (exon 2 to 5) | GTACAGACAGGAGGCGGATG | ACTCAAGGTCGCAGGTCAAG | cDNA amplifcation, sequencing |
| *Gfap* (ATG start - 3'UTR) | GAAGCAGGGCAAGATGGAGC | CCAGGCTGCTTGAACACAAC | cDNA cloning, sequencing |
| *Gfap* (3'UTR) | TCTGCCCTACCCACTCCTAC | | sequencing |
| *Gfap (* exon 6 to 8) | AACGTTAAGCTAGCCCTGGA | TTCCTCTTGAGGTGGCCTTC | qPCR, sequencing |
| *Lcn2** | CCGACACTGACTACGACCAG | AATGCATTGGTCGGTGGGAA | qPCR |
| *Cxcl10** | TGCAAGTCTATCCTGTCCGC | ACGGAGCTCTTTTTGACCTTC | qPCR |
| *Aif1** | AAGGATTTGCAGGGAGGAAAAGC | CTCCATGTACTTCGTCTTGAAGG | qPCR |
| *Rn18s* | CGCCGCTAGAGGTGAAATTCT | CGAACCTCCGACTTTCGTTCT | qPCR |

* reference (*65*)
